# Accurate Characterization of the Allosteric Energy Landscapes, Binding Hotspots and Long-Range Communications for KRAS Complexes with Effector Proteins : Integrative Approach Using Microsecond Molecular Dynamics, Deep Mutational Scanning of Binding Energetics and Allosteric Network Modeling

**DOI:** 10.1101/2025.01.27.635141

**Authors:** Sian Xiao, Mohammed Alshahrani, Guang Hu, Peng Tao, Gennady Verkhivker

**Affiliations:** Department of Chemistry, Center for Research Computing, Center for Drug Discovery, Design, and Delivery (CD4), Southern Methodist University, Dallas, Texas, 75275, United States of America; Keck Center for Science and Engineering, Schmid College of Science and Technology, Chapman University, Orange, CA 92866, United States of America; Department of Bioinformatics and Computational Biology, School of Life Sciences, Suzhou Medical College of Soochow University, Suzhou, 215213, China; Jiangsu Province Engineering Research Center of Precision Diagnostics and Therapeutics Development, Soochow University, Suzhou 215123, China; Department of Biomedical and Pharmaceutical Sciences, Chapman University School of Pharmacy, Irvine, CA 92618, United States of America

## Abstract

KRAS is a pivotal oncoprotein that regulates cell proliferation and survival through interactions with downstream effectors such as RAF1. Oncogenic mutations in KRAS, including G12V, G13D, and Q61R, drive constitutive activation and hyperactivation of signaling pathways, contributing to cancer progression. Despite significant advances in understanding KRAS biology, the structural and dynamic mechanisms of KRAS binding and allostery by which oncogenic mutations enhance KRAS-RAF1 binding and signaling remain incompletely understood. In this study, we employ microsecond molecular dynamics simulations, Markov State Modeling, mutational scanning and binding free energy calculations together with dynamic network modeling to elucidate the effect of KRAS mutations and characterize the thermodynamic and allosteric drivers and hotspots of KRAS binding and oncogenic activation. Our simulations revealed that oncogenic mutations stabilize the open active conformation of KRAS by differentially modulating the flexibility of the switch I and switch II regions, thereby enhancing RAF1 binding affinity. The G12V mutation rigidifies both switch I and switch II, locking KRAS in a stable, active state. In contrast, the G13D mutation moderately reduces switch I flexibility while increasing switch II dynamics, restoring a balance between stability and flexibility. The Q61R mutation induces a more complex conformational landscape, characterized by the increased switch II flexibility and expansion of functional macrostates, which promotes prolonged RAF1 binding and signaling. Mutational scanning of KRAS-RAF1 complexes identified key binding affinity hotspots, including Y40, E37, D38, and D33, and together with the MM-GBSA analysis revealed the hotspots leverage synergistic electrostatic and hydrophobic binding interactions in stabilizing the KRAS-RAF1 complexes. Network-based analysis of allosteric communication identifies critical KRAS residues (e.g., L6, E37, D57, R97) that mediate long-range interactions between the KRAS core and the RAF1 binding interface. The central β-sheet of KRAS emerges as a hub for transmitting conformational changes, linking distant functional sites and facilitating allosteric regulation. Strikingly, the predicted allosteric hotspots align with experimentally identified allosteric binding hotspots that define the energy landscape of KRAS allostery. This study highlights the power of integrating computational modeling with experimental data to unravel the complex dynamics of KRAS and its mutants. The identification of binding hotspots and allosteric communication routes offers new opportunities for developing targeted therapies to disrupt KRAS-RAF1 interactions and inhibit oncogenic signaling. Our results underscore the potential of computational approaches to guide the design of allosteric inhibitors and mutant-specific therapies for KRAS-driven cancers.

## Introduction

The GTPase KRAS (Kirsten rat sarcoma viral oncogene homolog) is a critical oncogene that is somatically mutated in approximately 10% of all human cancers, with particularly high prevalence in certain malignancies. Specifically, KRAS mutations are found in ∼90% of pancreatic adenocarcinomas, ∼40% of colorectal adenocarcinomas, ∼35% of lung adenocarcinomas, and ∼20% of multiple myeloma cases.^1–6^ Among the RAS oncogene family— which includes KRAS, HRAS, and NRAS—KRAS is the most frequently mutated, accounting for ∼30% of all RAS-driven cancers, while HRAS and NRAS mutations are less common, occurring in ∼8% and ∼3% of cases, respectively.^7,8^ These mutations drive tumorigenesis by disrupting normal cellular signaling pathways, leading to uncontrolled cell proliferation and survival.

RAS proteins function as binary molecular switches, cycling between an active GTP-bound state and an inactive GDP-bound state. This cycling is tightly regulated by two classes of proteins: Guanine nucleotide exchange factors (GEFs), which promote the exchange of GDP for GTP, activating RAS, and GTPase-activating proteins (GAPs), which enhance the intrinsic GTPase activity of RAS, facilitating GTP hydrolysis and returning RAS to its inactive state.^9^ Most oncogenic mutations in RAS proteins, particularly at residues G12, G13, and Q61, impair GTP hydrolysis and lock RAS in its active, GTP-bound conformation. These mutations introduce steric hindrance or alter the local structure of RAS, preventing GAPs from binding effectively.^10–13^ For example, mutations such as G12C, G12D, and Q61H alter the conformational dynamics of KRAS, leading to increased signaling activity and resistance to GAPs.^14–17^ As a result, RAS proteins remain constitutively active, leading to the overactivation of downstream signaling pathways such as the MAP kinase pathway (RAS-RAF-MEK-ERK) and the PI3K pathway (PI3K-AKT-mTOR). These pathways drive cell proliferation, survival, and metabolism, contributing to tumorigenesis.^18^

The interaction between KRAS and RAF1 (CRAF) is a critical step in the activation of the MAPK pathway, which regulates cell growth, differentiation, and survival. RAF1 is a serine/threonine kinase that binds to KRAS through two domains: the RAS-binding domain (RBD) and the cysteine-rich domain (CRD). Structural studies have provided detailed insights into the molecular mechanisms underlying this interaction. High-resolution crystal structures of KRAS bound to the RBD of RAF1 have revealed that the RBD interacts primarily with the switch I (residues 25–40) and switch II (residues 60–76) regions of KRAS.^19,20^ These regions undergo significant conformational changes during the GTPase cycle, transitioning between disordered (inactive) and ordered (active) states. The ordered conformation of switch I, stabilized by GTP binding, facilitates high-affinity binding to the RBD of RAF1, enabling downstream signaling. The CRD of RAF1, which is adjacent to the RBD, is thought to play a dual role in membrane association and allosteric regulation. The study presents high-resolution crystal structures of both wild-type and oncogenic mutants of KRAS in complex with the RBD and CRD of RAF1. These structures offer unprecedented insights into the molecular details of the KRAS-CRD interaction interface and reveal how the RBD and CRD are spatially organized relative to one another in the context of the KRAS-RAF1 complex. However, the precise mechanisms by which the CRD contributes to the stabilization of the active KRAS-RAF1 complex and facilitates signal transduction remain poorly understood. The N-terminal region of BRAF, including the CR1 and CR2 domains, remains unresolved in the absence of RAS^20^ which suggests that RAS binding is essential for stabilizing these domains and enabling proper RAF activation. Recent cryo-electron microscopy (cryo-EM) studies of full-length BRAF in complex with 14-3-3 and MEK have provided unprecedented views of RAF’s structural biology. ^21–25^ Oncogenic mutations in KRAS, such as G12V, G13D, and Q61R, do not disrupt the overall interaction with RAF1 but induce local rearrangements in the switch regions. X-ray crystallography of KRAS^Q61H^:GTP revealed that a hyperdynamic switch II allows for a more stable interaction with switch I, suggesting that enhanced RAF activity arises from a combination of absent intrinsic GTP hydrolysis activity and increased affinity for RAF.^26^ Similarly, the G13D mutation increases the flexibility of the P-loop, promoting the acquisition of active conformations in switch I and switch II, while the Q61R mutation disrupts the hydrogen bond network in the switch II region, increasing the conformational heterogeneity of KRAS.^10–12^ Over the past few decades, significant strides have been made in unraveling the structural intricacies of the RAS-RAF signaling pathway, a critical axis in cellular growth, differentiation, and survival. Despite these structural insights, the molecular basis of RAS-RAF binding— particularly the formation of the RAS-RBDCRD complex at the cell membrane—remained poorly understood.

The nanodisc platforms and paramagnetic relaxation enhancement (PRE) analyses were employed to determine the structure of a hetero-tetrameric complex comprising KRAS and the RBD and CRD of activated RAF1.^25^ By leveraging these advanced techniques, this study investigated how the binding of the RBD or the RBD–CRD differentially modulates the dimerization modes of KRAS on both anionic and neutral membranes. A key discovery of this work is that RBD binding allosterically generates two distinct KRAS dimer interfaces, which exist in a dynamic equilibrium. One interface is favored when KRAS is free, while the other is stabilized when KRAS is in complex with the RBD–CRD.^25^ This allosteric regulation highlights the plasticity of KRAS dimerization and its responsiveness to effector binding. Biochemical and structural analyses of variants identified in a KRAS^G12D^ revealed that attenuation of oncogenic KRAS can be mediated by protein instability and conformational rigidity, resulting in reduced binding affinity to effector proteins or reduced SOS-mediated nucleotide exchange activity. These studies define the landscape of single amino acid alterations that modulate the function of KRAS.^27^

KRAS has long been considered “undruggable” due to its smooth surface and lack of deep, well-defined binding pockets. However, recent advances in structural biology identified cryptic allosteric pockets that can be targeted by small molecules. These pockets are transiently formed during conformational changes and play a critical role in regulating KRAS activity through allosteric mechanisms. For example, the approval of sotorasib (AMG 510), a covalent inhibitor that targets the KRAS(G12C) mutation, binds to a cryptic pocket outside the nucleotide-binding and effector-binding sites, locking KRAS(G12C) in its inactive, GDP-bound state and preventing interaction with downstream effectors.^16^ A group of monobodies, small synthetic binding proteins was developed that were selective to KRAS(G12D) over KRAS wild type (WT) and other oncogenic KRAS mutations.^28^ Crystallographic studies revealed that, similar to other KRAS mutant-selective inhibitors, the initial monobody bound to the S-II pocket, the groove between switch II and α3 helix, and captured this pocket in the most widely open known form.^28^

A sophisticated comprehensive analysis of KRAS energetics and allostery was undertaken using a combination of structural biology and biophysical techniques to map the energetic and allosteric landscape of KRAS.^29^ This seminal study characterized allosteric energy landscapes that highlighted the thermodynamic consequences of mutations across KRAS providing a quantitative framework to determine allosteric binding sites and their effect on KRAS binding. The use of double mutants allowed to probe genetic interactions and infer causal free energy changes at scale providing a high-resolution view of how mutations influence KRAS folding and binding.^29^ By analyzing the free energy changes associated with each mutation, the residues and regions that are critical for maintaining KRAS stability and binding specificity were determined. The study reveals the existence of extensive allosteric networks within KRAS that connect distant regions of the protein. These networks facilitate communication between the nucleotide-binding site, effector-binding site, and other regulatory regions. The authors also identified several cryptic pockets on the surface of KRAS that are transiently formed during conformational changes.^29^ These pockets can be targeted by small molecules to modulate KRAS activity.

While experimental techniques like crystallography and cryo-EM have provided valuable structural insights, computational studies and molecular modeling have played an equally important role in elucidating the dynamics, energetics, and molecular mechanisms of KRAS-RAF1 binding. MD simulations, free energy calculations, and docking studies have complemented experimental techniques, revealing the role of conformational flexibility, membrane interactions, and allosteric regulation. MD simulations have revealed that the switch I and switch II regions of KRAS are highly dynamic, adopting multiple conformations that influence its binding to RAF1. MD simulations indicated similarities in conformational mobility of KRAS-WT and mutants, thus revealing complexity of mechanisms in which oncogenic mutations (e.g., G12C, G12D) stabilize the active conformation of KRAS, enhancing its affinity for RAF1. MD simulations performed by Lu et al. suggested that conformational transformation of the switch domain because of Q61H affects intrinsic GTPase activity.^11^ MD simulations also suggested that simulations that Q61H mutation in the switch-II does not generate clear effect on the fluctuations of the switch-II in active KRAS but may modulate the flexibility of inactive KRAS protein.^31^ Multiple replica Gaussian accelerated molecular dynamics (MR-GaMD) simulations were applied to decode the effect of Q61A, Q61H and Q61L on the activity of KRAS revealing that dynamics behavior of the switch domain in KRAS is affected by the three Q61 mutants, inducing structural flexibility of the switch domain and disturb the activity of KRAS.^12^ Multiscale coarse-grained and all-atom MD simulations of KRAS4b bound to the RBD and CRD domains of RAF-1, both in solution and anchored to a model plasma membrane explored how RAS membrane orientation relates to the protein dynamics within the RAS-RBDCRD complex.^32^ Solution MD simulations describe dynamic KRAS4b-CRD conformations, suggesting that the CRD has sufficient flexibility in this environment to substantially change its binding interface with KRAS4b. Quantitative spatial imaging and atomistic molecular dynamics simulations to examine molecular details of K-Ras plasma membrane binding.^33^ The effects of amino acid variations on the structure and dynamics of KRAS-WT and oncogenic mutants G12D, G12V, and G13D of HRAS and KRAS proteins. Based on data from µs-scale molecular dynamics simulations, we show that the overall structure of the proteins remains similar but there are important differences in dynamics and interaction networks.^34^ The insightful study employed all-atom molecular dynamics simulations of the C-RAF RBD and CRD regions when bound to oncogenic KRAS4B suggesting that the membrane plays an integral role in regulating the configurational ensemble of the complex that samples a few states dynamically, reflecting a competition between CRAF CRD- and KRAS4B-membrane interactions.^35^ Microsecond MD simulations to unveil the binding mechanisms of the FDA-approved MEK inhibitor trametinib with KRASG12D, providing insights for potential drug repurposing. The binding of trametinib was compared with clinical trial drug MRTX1133, which demonstrates exceptional activity against KRASG12D, for better understanding of interaction mechanism of trametinib with KRASG12D.^36^ Multiple microsecond MD simulations and Markov State Model (MSM) analysis probed kinetics of MRTX1133 binding to KRAS G12D revealing the kinetically metastable states and potential pathways of MRTX1133 binding while MM/GBSA analysis identified 8 critical residues for MRTX1133 recognition and binding.^37^ KRAS binding pockets and ligand interactions, identifying three distinct sites: the conserved nucleotide-binding site, the shallow switch-I/II pocket, and the allosteric Switch-II/α3 pocket.^38^ In addition, this study identified KRAS-inhibitor interaction fingerprints aided by MD simulations that characterized the flexibility of these sites to accommodate diverse ligands.^38^ MD simulations of the KRASG12C-AMG 510 complex investigated the impact of ligand binding on KRASG12C conformational changes.^39^ Mapping simulation trajectories onto the Principal Component Analysis (PCA) model revealed that KRASG12C-AMG 510 predominantly adopted the inactive conformation where AMG 510 reduced the flexibility of the switch regions, stabilizing the KRASG12C-AMG 510 complex in the inactive state.^39^ MD simulations explored the structural dynamics and stability of wild-type KRAS and its oncogenic variants (G12C, G12D, G12V, G13D) followed by the free energy landscape analysis revealing unique conformational dynamics and altered thermodynamic stability in mutated KRAS variants.^40^ Multiple MD simulations were performed on wild-type and mutant KRAS structures to investigate how G12C and G12D mutations stabilize the active state and how AMG-510 and MRTX1133 inhibitors force these mutants into the inactive state.^41^ The study revealed that binding of AMG-510 (Sotorasib) can induce significant stabilization of the Switch-II region of KRAS-G12C, surpassing that of MRTX1133 bound with KRAS-G12D mutant. Mutation probabilities of KRAS-G12 missense mutants and their long-timescale dynamics were assessed by atomistic MD simulations (170 μs) and MSM analysis revealing allosteric hydrophobic signaling network in KRAS, and allosteric modulation of protein dynamics among the G12X mutants which is manifested in the switch regions that are responsible for the effector protein binding.^42^ GaMD simulations were performed on the GDP-bound wild-type, G12A, G12D, and G12R KRAS to probe mutation-mediated impacts on conformational alterations of KRAS, showing that all three G12 mutations can alter the structural flexibility of the switch domains.^43^ The analyses of the free energy landscapes indicated that the examined G12 mutations can induce more conformational states of KRAS and particularly result in more disordered switch domains.^43^ GaMD) simulations followed by deep learning (DL) were carried out to probe the effect of G12C mutation and binding of three proteins NF1, RAF1, and SOS1 on the conformational dynamics and free energy landscapes of KRAS4B, showing that mutation and binding alter contacts in key structural domains and enhance switch I and switch II mobility.^44^ These findings highlighted the roles of partner binding and G12C in KRAS4B activity and allosteric regulation, providing theoretical insights into KRAS4B function. GaMD simulations and DL also explored the phosphorylation-mediated effect on conformational dynamics of the GTP-bound KRAS revealing that the phosphorylation of pY32, pY64, and pY137 sites leads to more disordered states of the switch domains and can induce conformational transformations between the closed and open states.^45^

To summarize, computational studies of KRAS revealed several key observations. First, MD simulations highlighted the high flexibility of the Switch I and Switch II regions in KRAS, which adopt multiple conformations influencing RAF1 binding. Second, Oncogenic mutations (e.g., G12C, G12D) stabilize the active conformation of KRAS, enhancing its affinity for RAF1, while maintaining similarities in conformational mobility with wild-type KRAS. In particular, Q61 mutations (e.g., Q61H, Q61A, Q61L) affect the intrinsic GTPase activity and structural flexibility of the switch domains, particularly in inactive KRAS. Multiscale MD simulations also suggested that KRAS membrane orientation and interactions with RAF1 RBD/CRD domains are critical for protein dynamics and complex formation. The analysis of mutation-specific effects showed that G12 mutations (e.g., G12C, G12D, G12V, G12R) induce conformational changes, particularly in the switch domains, leading to disordered states and altered thermodynamic stability. While G12C destabilizes key structural domains (switch I, switch II, loop 4, and loop 5), protein binding partners (e.g., NF1, RAF1, SOS1) stabilizes these regions, highlighting the interplay between mutations and partner interactions. Computational studies of inhibitor binding also showed that AMG-510 (Sotorasib) and MRTX1133 lock KRAS mutants (G12C, G12D) in the inactive state by stabilizing the Switch-II region, with AMG-510 showing greater stabilization for G12C.

Despite the significant body of structural, biochemical and computational studies of KRAS dynamics and binding, there are a number of open questions from understanding the fundamental biophysics of KRAS dynamics and allosteric networks to addressing practical challenges in allosteric drug design and resistance. The important open questions include the following issues : (a) How do long-timescale conformational changes in KRAS, particularly in the switch regions influence its binding to effectors (e.g., RAF1, PI3K, RALGDS) and regulators (e.g., GAPs, GEFs)? (b) What are allosteric networks that connect distal mutation sites (e.g., G12, Q61) to the switch regions, and how do these networks modulate KRAS activity? (c) What are the structural and dynamic differences between various KRAS mutations (e.g., G12C, G12D, G12V, G13D, Q61H) that lead to oncogenic outcomes? (d) How do KRAS mutations alter the free energy landscape of conformational states, and what are the thermodynamic drivers of mutation-induced oncogenic activation? (e) What are the energetic contributions of specific residues and interactions to KRAS stability, and allosteric effector binding?

In this study, we attempted to address some of these issues by employing an integrative computational simulation strategy that employed multiple microsecond MD simulations of KRAS-RAF1 and KRAS oncogenic mutants G12V, G13D and Q61R in the complex with RBD of RAF1, MSM analysis, systematic “deep” mutational scanning (DMS) of KRAS residues for binding and stability, rigorous MM-GBSA binding free energy analysis and network modeling of allosteric networks to determine binding affinity hotspots and allosteric binding centers. In addition, for structure-based mutational scanning we also employed the crystal structures of KRAS-Q61R (GMPPNP-bound) and KRAS-G13D (GMPPNP-bound) in complex with RBD and CRD of RAF1/CRAF. These simulations provide a detailed understanding of conformational flexibility and allosteric communication within the KRAS-RAF1 complex, revealing how oncogenic mutations alter the dynamics of KRAS and enhance its interaction with RAF1. The results of simulations, binding analysis and allosteric network modeling performed in this study are compared in detail with the pioneering experimental study by Lehner and colleagues that characterized the energetic and allosteric landscape for KRAS inhibition.^29^ The ensemble-based mutational profiling of KRAS residues using knowledge-based energy model enabled accurate identification of binding affinity and protein stability hotspots. The detailed MM-GBSA analysis of binding energetics reveals the thermodynamic drivers of binding and the energetic contributions of the binding affinity hotspots to KRAS stability and effector binding, showing excellent agreement with the experimental DMS data.^29^ Using MD-inferred conformational ensembles and dynamics-based network modeling, this study mapped potential allosteric hotspots and allosteric communication pathways in KRAS-RAF1 complexes. Consistent with rigorous experimental analysis, our results confirmed that the central β-sheet of KRAS acts as a hub for transmitting allosteric signal between distant functional sites, facilitating allosteric communication between the switch regions and the RAF1 binding interface. Moreover, network-based dynamic modeling of allostery predicted key allosteric hotspots in which mutations or structural perturbations can dramatically affect the fidelity of allosteric communication. Strikingly, these predictions almost precisely reproduced the experimental data on allosteric sites in which mutations have strong allosteric effects on the binding free energy with RAF1 that is equal to or greater than that in binding interface residues.^29^ The vast majority of 18 experimentally detected allosteric binding hotspots were predicted in the network analysis of allostery that also provided a detailed map of allosteric pathways between the allosteric centers and binding interface hotspots in the KRAS-RAF1 complex.

The results of this study provide important quantitative insights into principles of allosteric communication and allosteric binding in KRAS that are consistent with and can explain the experimental data. In particular, we find that allosteric communication hotspots are enriched close to the functional switch regions and leverage conformational plasticity in these regions for transmitting allosteric signals, also suggesting that local energetic propagation as the main allosteric mechanism. The results also confirm that allosteric communication is anisotropic and is mediated by conserved positions that is particularly effective across the central β-sheet of KRAS.

## Materials and Methods

### Structural Analysis and Preparation of Simulation Systems

The crystal and cryo-EM structures of the KRAS and KRAS complexes are obtained from the Protein Data Bank.^46^ Microsecond MD simulations are performed for the crystal structure of wild-type KRAS4b (GMPPNP-bound) in complex with the RBD of RAF1/CRAF (pdb id 6VJJ); G12V KRAS mutant in the complex with the RBD of RAF1/CRAF (pdb id 6VJJ); crystal structure of KRAS-G13D (GMPPNP-bound) in complex with the RBD and CRD of RAF1/CRAF (pdb id 6XGV); the crystal structure of KRAS-Q61R (GMPPNP-bound) in complex with RBD and CRD of RAF1/CRAF (pdb id 6XGU). For simulated structures, hydrogen atoms and missing residues were initially added and assigned according to the WHATIF program web interface.^47^ Missing residues or loops were modeled using MODELLER^48^, and the structures were cleaned to remove water molecules, ligands, and non-standard residues. The tleap module in the AMBER package was also employed to add the missing hydrogen atoms of the structures.^49^ The missing regions are subsequently reconstructed and optimized using template-based loop prediction approach ArchPRED.^50^ The side chain rotamers were refined and optimized by SCWRL4 tool.^51^ Protonation states of ionizable residues were assigned using PROPKA, ensuring histidine residues were in the appropriate tautomeric state.^52,53^ The tleap module in AMBER was used to generate the topology (prmtop) and coordinate (inpcrd) files. The system was solvated in a TIP3P water box with a 12 Å buffer and neutralized using Na+ or Cl-ions. The final system was saved for further simulations. Molecular mechanics parameters of proteins were assigned according to the ff14SB force field.^54,55^ The protein structures were energy-minimized to remove steric clashes and optimize the structure. Two stages of minimization were performed: (1) with restraints on the protein backbone to relax solvent and ions, and (2) without restraints to minimize the entire system. The minimization was conducted using the pmemd.cuda module with the following parameters: 1,000 cycles of steepest descent followed by 500 cycles of conjugate gradient minimization, a cutoff of 10 Å for non-bonded interactions, and the ff14SB force field for proteins.^54,55^ The parameters for the ligands are generated by Antechamber using GAFF2 force field.^56,57^

### All-Atom Molecular Dynamics Simulations

The relaxation process includes minimization, heating, restriction run, and equilibrium run. After minimization, the system is heated from 100K to 300K, over 1 nanosecond of simulation time at a constant volume, with integration time 1 fs. Then we relax system at a constant pressure with protein restraints over 1 ns of simulation time at a constant pressure with restraints set to 1 kcal/mol Å^2. Finally, we relax the system with no restraints for 1 ns of simulation time at a constant pressure. Production MD simulations were performed for 2µs under NPT conditions (300 K, 1 bar) using the pmemd.cuda module. The temperature was maintained using the Langevin thermostat, and pressure was controlled with the Monte Carlo barostat. A time step of 2 fs was used, and bonds involving hydrogen atoms were constrained using the SHAKE algorithm. MD trajectories were saved every 100 ps for analysis. The ff14SB force field was used for proteins, and the TIP3P model was used for water. For each system, MD simulations were conducted three times in parallel to obtain comprehensive sampling. Each individual simulation has 10,000 frames.

The CPPTRAJ software in AMBER 18 was used to calculate the root mean squared deviation (RMSD) and root mean squared fluctuation (RMSF) of MD simulation trajectories in which the initial structure was used as the reference.^58^ The structures of were visualized using Visual Molecular Dynamics (VMD 1.9.3) ^59^ and PyMOL (Schrodinger, LLC. 2010. The PyMOL Molecular Graphics System, Version X.X.)

To identify the dynamical coupling of the motions between protein segments, the cross-correlation coefficient (*C_ij_*) was proposed for measuring the motion correlation between the C_α_ atom pair in residues *i* and *j*, which is defined as

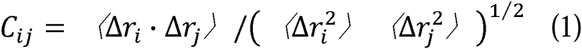

where and are the displacement from the mean position of the C_α_ atom pair in residues *i* and *j*, considered over the sampled period. Positive *C_ij_* denotes lockstep motion in the residue pair under investigation, whereas negative *C_ij_* stands for negatively correlated motion. The cross-correlation analysis in this work was implemented using the Bio3D package.^60^ This methodology provides a comprehensive framework for studying KRAS-RAF1 binding dynamics using AMBER, enabling detailed insights into the conformational changes and interactions governing this critical protein-protein interaction.

### Markov State Model

Stochastic Markov state models (MSMs)^61–63^ have become increasingly useful states-and-rates models with the mature and robust software infrastructure^64,65^ for describing the transitions between functional protein states and modeling of allosteric events. In MSM, protein dynamics is modeled as a kinetic process consisting of a series of Markovian transitions between different conformational states at discrete time intervals. A specific time interval, referred to as lag time, needs to be determined to construct transition matrix. First, k-means clustering method is conducted on projected low-dimensional space and each simulation frame is assigned to a microstate. The transition counting is constructed based on a specific time interval lag time τ. Macrostates are kinetically clustered based on the Perron-cluster cluster analysis (PCCA++)^66^ and considered to be kinetically separate equilibrium states. The transition matrix and transition probability were calculated to quantify the transition dynamics among macrostates. The corresponding transition probability from state *i* to state *j* is calculated as:

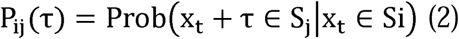

A proper lag time is required for MSM to be Markovian. The value of the lag time and the number of macrostates are selected based on the result of estimated relaxation timescale.^67^ The implied timescales can be calculated using the eigenvalues (*λ_i_*) in the transition matrix as

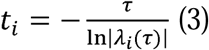

The number of protein metastable states associated with these slow relaxation timescales can be inferred based on the convergence of implied relaxation time scale. These metastable states effectively discretize the conformational landscape. The MSM building was conducted using PyEMMA package (v2.5.12).^68^ Based on the transition matrix we obtain implied timescales for transitions between various regions of phase space and use this information determines the number of metastable states. The number of metastable states also defines the resolution of the model by determining how large a barrier must be in order to divide phase space into multiple states.

### Ensemble-Based Mutational Scanning and Sensitivity Analysis of the KRAS Residues : Quantifying Effects of Mutations on KRAS Binding and Protein Stability

We conducted a systematic mutational scanning analysis of the KRAS residues in the KRAS complexes using conformational ensembles of KRAS-RAF1 complexes and averaging of mutation-induced energy changes. Every KRAS residue was systematically mutated using all substitutions and corresponding protein stability and binding free energy changes were computed with the knowledge-based BeAtMuSiC approach.^69–71^ This approach is based on statistical potentials describing the pairwise inter-residue distances, backbone torsion angles and solvent accessibilities, and considers the effect of the mutation on the strength of the interactions at the interface and on the overall stability of the complex. The binding free energy of protein-protein complex can be expressed as the difference in the folding free energy of the complex and folding free energies of the two protein binding partners:

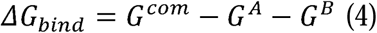

The change of the binding energy due to a mutation was calculated then as the following:

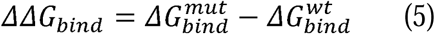

We leveraged rapid calculations based on statistical potentials to compute the ensemble-averaged binding free energy changes using equilibrium samples from simulation trajectories. The binding free energy changes were obtained by averaging the results over 1,000 and 10, 000 equilibrium samples for each of the systems studied.

### MM-GBSA Binding Free Energy Computations of KRAS-RAF1 Complexes

We calculated the ensemble-averaged changes in binding free energy using 1,000 equilibrium samples obtained from simulation trajectories for each system under study. The binding free energies of the KRJAS-RAF1 complexes were assessed using the MM-GBSA approach.^72,73^ The energy decomposition analysis evaluates the contribution of each amino acid to binding of KRAS to RAF1 protein.^74,75^ The binding free energy for the KRAS-RAF1 complex was obtained using:

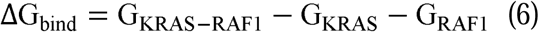

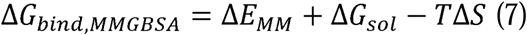

where Δ*E_MM_* is total gas phase energy (sum of Δ*E_internal_*, Δ*E_electrostatic_*, and Δ*Evdw*); Δ*Gsol* is sum of polar (Δ*G_GB_*) and non-polar (Δ*G_SA_*) contributions to solvation. Here, G_KRAS–RAF1_ represent the average over the snapshots of a single trajectory of the complex, G_KRAS_ and G_RAF1_ corresponds to the free energy of KRAS and RAF1 protein, respectively. The polar and non-polar contributions to the solvation free energy is calculated using a Generalized Born solvent model and consideration of the solvent accessible surface area.^76^ MM-GBSA is employed to predict the binding free energy and decompose the free energy contributions to the binding free energy of a protein–protein complex on per-residue basis. The binding free energy with MM-GBSA was computed by averaging the results of computations over 1,000 samples from the equilibrium ensembles. First, the computational protocol must be selected between the “single-trajectory” (one trajectory of the complex), or “separate-trajectory” (three separate trajectories of the complex, receptor and ligand). To reduce the noise in the calculations, it is common that each term is evaluated on frames from the trajectory of the bound complex. In this study, we choose the “single-trajectory” protocol, because it is less noisy due to the cancellation of intermolecular energy contributions. This protocol applies to cases where significant structural changes upon binding are not expected. Entropy calculations typically dominate the computational cost of the MM-GBSA estimates. Therefore, it may be calculated only for a subset of the snapshots, or this term can be omitted.^77,78^ However, for the absolute affinities, the entropy term is needed, owing to the loss of translational and rotational freedom when the ligand binds. In this study, the entropy contribution was not included in the calculations of binding free energies of the complexes because the entropic differences in estimates of relative binding affinities are expected to be small owing to small mutational changes and preservation of the conformational dynamics.^77,78^ MM-GBSA energies were evaluated with the MMPBSA.py script in the AmberTools21 package.^79^

### Graph-Based Dynamic Network Analysis of Protein Ensembles

To analyze protein structures, we employed a graph-based representation where residues are modeled as network nodes, and non-covalent interactions between residue side-chains define the edges. This approach captures the spatial and functional relationships between residues, providing insights into the protein’s structural and dynamic properties. The graph-based framework allows for the integration of both structural and evolutionary information, enabling a comprehensive analysis of residue interactions. The residue interaction networks were constructed by defining edges based on non-covalent interactions between residue side-chains.^80,81^ The weights of these edges were determined using two key metrics: (a) dynamic residue cross-correlations derived from MD simulations where these correlations quantify the coordinated motions of residue pairs^82^; and (b) coevolutionary couplings measured using mutual information scores, where these couplings reflect evolutionary constraints and residue-residue dependencies.^83^ The edge lengths (weights) between nodes ii and jj were computed using generalized correlation coefficients, which integrate both dynamic correlations and coevolutionary mutual information. Only residue pairs observed in at least one independent simulation were included in the network. The matrix of communication distances was constructed using generalized correlations between composite variables that describe both the dynamic positions of residues and their coevolutionary mutual information. This matrix provides a quantitative measure of communication efficiency between residues, reflecting both their physical proximity and evolutionary relationships. The Residue Interaction Network Generator (RING) program^84–87^ was used to generate residue interaction networks from the conformational ensemble. The edges in these networks were weighted to reflect the frequency of interactions observed in the ensemble. Network files in XML format were generated for all structures using the RING v3.0 webserver. Network graph calculations were performed using the Python package NetworkX.^88,89^ This included the computation of key network parameters, such as shortest paths and betweenness centrality, to identify residues critical for communication within the protein structure. The short path betweenness (SPC) of residue *i* is defined to be the sum of the fraction of shortest paths between all pairs of residues that pass through residue *i*:

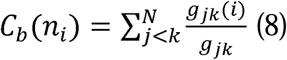

where *g_jk_* denotes the number of shortest geodesics paths connecting *j* and *k,* and *g _jk_* (*i*) is the number of shortest paths between residues *j* and *k* passing through the node *n_i_.* Residues with high occurrence in the shortest paths connecting all residue pairs have a higher betweenness values. For each node *n,* the betweenness value is normalized by the number of node pairs excluding *n* given as(*N* −1)( *N* - 2) / 2, where *N* is the total number of nodes in the connected component that node *n* belongs to. To account for differences in network size, the betweenness centrality of each residue ii was normalized by the number of node pairs excluding ii. The normalized short path betweenness of residue *i* can be expressed as :

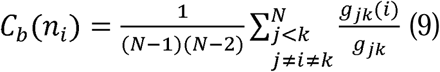

*g_jk_* is the number of shortest paths between residues *j* and k; *g_jk_* (*i*) is the fraction of these shortest paths that pass through residue *i*.

Residues with high normalized betweenness centrality values were identified as key mediators of communication within the protein structure network. These residues are likely to play critical roles in maintaining the protein’s structural integrity and functional dynamics.

### Network-Based Mutational Profiling of Allosteric Residue Centrality

Through mutation-based perturbations of protein residues we compute dynamic couplings of residues and changes in the short path betweenness centrality (SPC) averaged over all possible modifications in a given position.

The change of SPC upon mutational changes of each node is reminiscent to the calculation of residue centralities by systematically removing nodes from the network.

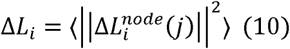

where *i* is a given site, *j* is a mutation and 〈⋯〉 denotes averaging over mutations. 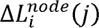 describes the change of SPC parameters upon mutation *j* in a residue node *i*. Δ*L_i_* is the average change of ASPL triggered by mutational changes in position *i*.

Z-score is then calculated for each node as follows^108,109^:

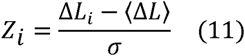

〈Δ*L*〉 is the change of the SPC network parameter under mutational scanning averaged over all protein residues and σ is the corresponding standard deviation. The ensemble-average Z score changes are computed from network analysis of the conformational ensembles of KRAS-RAF1 complexes using 1,000 snapshots of the simulation trajectory. Through this approach, we evaluate the effect of mutations in the KRAS residues on allosteric communications with RAF1.

## Results

### Microsecond MD Simulations of the KRAS-WT, G12V, G13D and Q61R Complexes with RAF1 Reveal Distinct Dynamic Signatures of RBD Proteins

MD simulations of the crystal structure of KRAS4b WT (GMPPNP-bound) in complex with the RBD of RAF1 (pdb id 6VJJ) provided detailed key insights into the dynamic behavior, stability, and interactions of this complex. A complete GTPase reaction requires well-ordered conformations of the protein active site, which includes the phosphate-binding loop, P-loop (residues 10–17), switch I (residues 25–40) and switch II (residues 60–76) regions (Figure 1).

**Figure 1.**
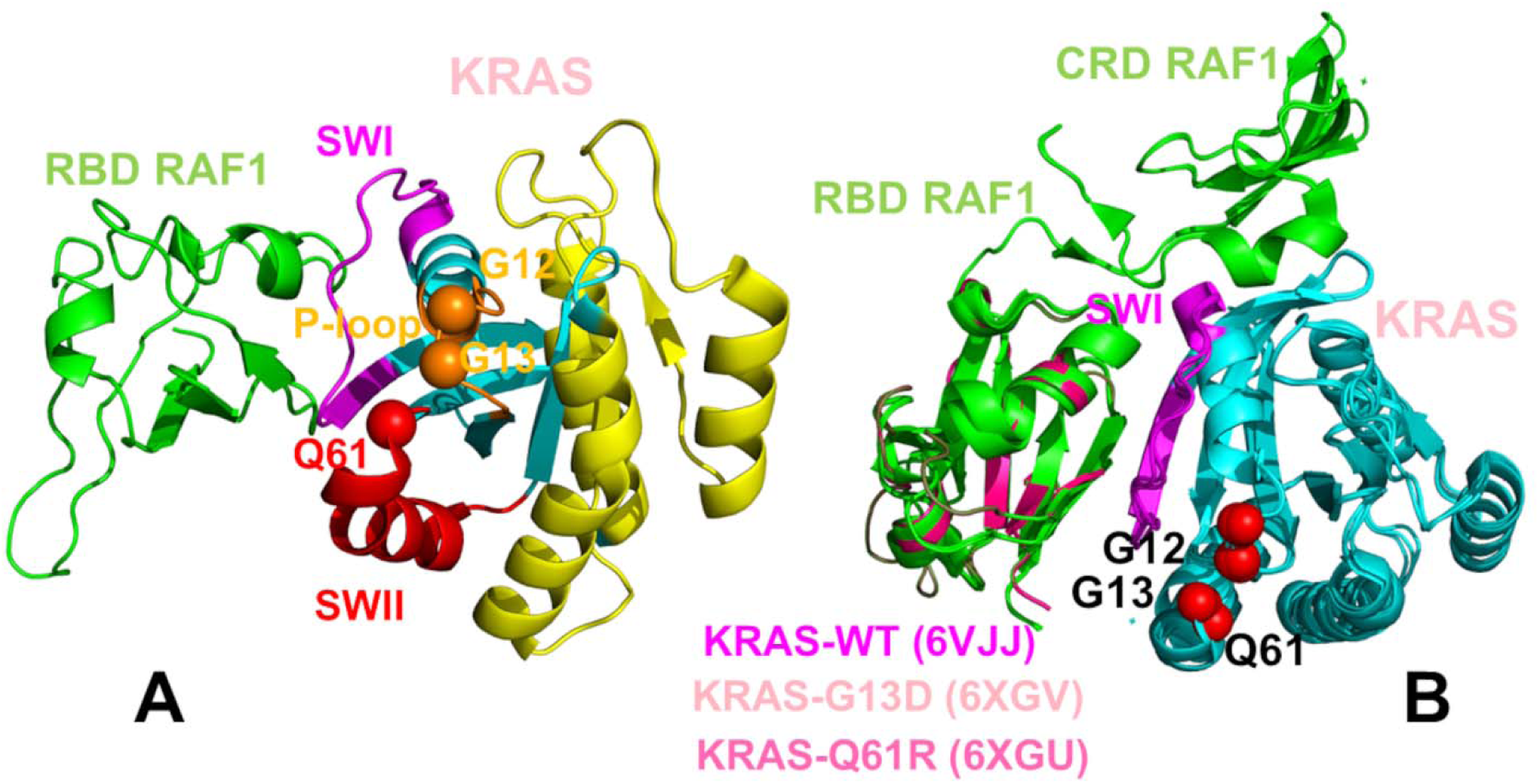
Structural overview and organization of the KRAS protein with RAF1. (A) The crystal structure of wild-type KRAS (GMPPNP-bound) in complex with the RBD of RAF1/CRAF (pdb id 6VJJ). The functional KRAS regions are highlighted P-loop (residues 10-17 in orange ribbons), : switch I, SW1 (residues 24-40 in magenta ribbons), switch II, SWII (residues 60-76 in red ribbons), allosteric KRAS lobe (residues 87-166 in yellow ribbons). The sites of oncogenic mutants G12V, G13D and Q61R are shown in spheres colored according to the color of the respective KRAS functional region to which they belong. (B) Structural overlay of the crystal structure of wild-type KRAS (GMPPNP-bound) in complex with the RBD of RAF1/CRAF (pdb id 6VJJ), (crystal structure of KRAS-G13D (GMPPNP-bound) in complex with the RBD and CRD of RAF1/CRAF (pdb id 6XGV) and the crystal structure of KRAS-Q61R (GMPPNP-bound) in complex with RBD and CRD of RAF1/CRAF (pdb id 6XGU). The sites of oncogenic mutants G12V, G13D and Q61R are shown in red spheres.

The conformational flexibility of the KRAS and RAF1 was analyzed by calculating the root mean square deviations (RMSD) (Supplementary Materials, Figure S1) and the root mean square fluctuations (RMSF) distribution for the KRAS residues in different complexes (Figure 2). The RMSD profiles for the KRAS residues demonstrated consistent convergence of the MD trajectories for the KRAS-WT complex for all three trajectories reaching a stable equilibrium state after approximately 500 ns (Supplementary Materials, Figure S1A). A synchronous behavior was seen for three independent microsecond MD trajectories for KRAS-G12V where full convergence is typically reached after 800 ns – 1 µs (Supplementary Materials, Figure S1B). However, a noticeable divergence in RMSD profiles was observed for the MD trajectories of the KRAS-G13D complex (Supplementary Materials, Figure S1C), reflecting a more heterogeneous ensemble of KRAS conformations in this variant. For the KRAS-Q61R complex, the RMSD profiles showed considerably more heterogeneity among three microsecond trajectories, stabilizing after ∼500-700 ns (Supplementary Materials, Figure S1D). The RMSDs for RAF1 also displayed variability across trajectories for the KRAS-WT, G13D, and Q61R variants, highlighting the functionally significant plasticity of both binding partners in the KRAS-RAF1 complexes (Supplementary Materials, Figure S2). These preliminary findings suggest a certain degree of dynamic cooperativity between KRAS and RBD RAF1 proteins in response to oncogenic mutation where the increased plasticity of KRAS in the Q61R mutant can induce the reciprocal increase in thermal fluctuations for the interacting RBD RAF1 (Supplementary Materials, Figure S2D) compared to the smaller deviations between RAF1 trajectories for KRAS-WT (Supplementary Materials, Figure S2A) and KRAS-G12V complexes (Supplementary Materials, Figure S2B).

**Figure 2.**
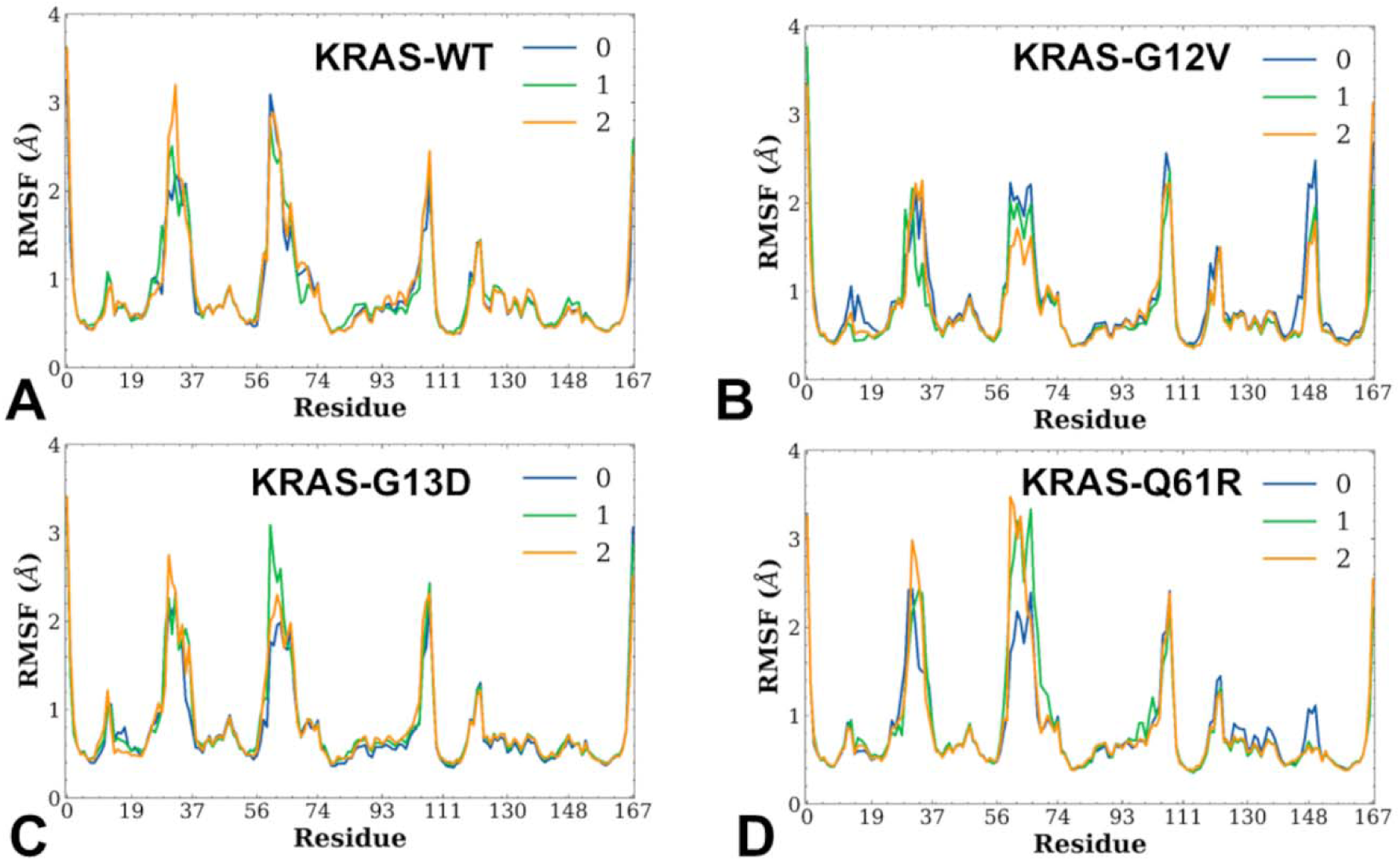
Conformational dynamics profiles of the KRAS protein residues obtained from three independent MD simulations of the KRAS complexes with RAF1. The RMSF profiles for the RBD residues obtained from three independent 2 µs MD simulations of the crystal structure of wild-type KRAS (GMPPNP-bound) in complex with the RBD of RAF1/CRAF (pdb id 6VJJ); (A), KRAS-G12V mutant in the complex with the RBD of RAF1/CRAF (pdb id 6VJJ); (B), crystal structure of KRAS-G13D (GMPPNP-bound) in complex with the RBD and CRD of RAF1/CRAF (pdb id 6XGV) (C) and the crystal structure of KRAS-Q61R (GMPPNP-bound) in complex with RBD and CRD of RAF1/CRAF (pdb id 6XGU) (D).

The RMSF profiles revealed that the KRAS4b-RAF1 WT complex is stable over the simulation timescale, with the RBD of RAF1 maintaining its binding to the switch I and switch II regions of KRAS4b (Figure 2A). These regions are critical for GTP binding, effector interaction, and downstream signaling. The P-loop is a highly conserved region in KRAS that interacts with the phosphate groups of GTP. It plays a critical role in nucleotide binding and stabilization. MD simulations showed that the P-loop (residues 10-17) exhibits moderate flexibility that allows the P-loop to adapt to the binding of GTP and accommodate conformational changes during the GTPase cycle (Figure 2A). The switch I region displayed considerable flexibility, particularly evident in one of the three independent microsecond simulations, fluctuating between short-lived partly disordered/semi-open (inactive) states and dominant ensemble of ordered/open (active) states (Figure 2A). According to structural studies, in the GDP-bound state, switch I is largely disordered, while in the GTP-bound state it adopts an ordered conformation that facilitates effector binding. MD simulations suggested that in the KRAS-RAF1 RBD complex, the switch I region of KRAS can adopt several semi-open conformational states with the increased mobility, but the population is dominated by the fully ordered open conformations (Figure 2A). A small fraction of the detected semi-open conformations are likely intermediate states between the closed (inactive) and open (active) conformations of switch I. It is possible that this transient form may occur during the transition from the GDP-bound to the GTP-bound form or during the initial stages of effector binding. On the other hand, the prevalent open conformation of switch I is the fully active state that is stabilized by GTP binding and enables high-affinity binding to the RBD of RAF1. The critical residues in the switch I region, such as Y32, D33 and E37 sites^20^ are fully exposed and available for interaction with the RBD of RAF1 in the dominant open state (Figure 2A) where particularly D33 forms a salt bridge with R59 in the RBD and E37 interacts with R67 in the RBD.

The G12V mutation in KRAS is one of the most common oncogenic mutations and has effects on the structure, dynamics, and function of KRAS, particularly in its interaction with the downstream effector RAF1. We employed the KRAS-WT structure in the complex with RAF1 (pdb id 6VJJ) as a template for introducing G12V mutation followed minimization and equilibration of the resulting complex. Conformational dynamics profiles for KRAS-G12V complex clearly indicated as can be judged from three independent microsecond simulations that the G12V mutation can reduce flexibility of the switch I and II regions thus stabilizing the open conformation. Our results may seem counterintuitive at first, as oncogenic mutations like G12V are often associated with increased flexibility in these regions. However, this observation can be explained by considering the specific structural and dynamic effects of the G12V mutation, as well as the context of the simulations (Figure 2B). The switch I region is directly connected to the P-loop and is sensitive to changes in the nucleotide-binding site (Figure 1). According to our results, the G12V mutation may restrict the flexibility of switch I region by stabilizing its interactions with GTP and the P-loop (Figure 2B). Switch II region is also influenced by the P-loop and the nucleotide-binding site. The G12V mutation may reduce switch II fluctuations by stabilizing the overall active conformation of KRAS, including the orientation of switch II relative to switch I and the nucleotide. This stabilization may reduce the conformational entropy of the switch regions by favoring a specific, stable open state favorable for RAF1 effector binding over a dynamic ensemble of states (Figure 2B). Through these dynamic changes oncogenic G12V mutation may promote prolonged effector binding contributing to oncogenic transformation. In addition, the reduced flexibility of switch I limits its ability to adopt alternative conformations, which may affect its interaction with other regulatory proteins.

Structural mapping of MD snapshots for the KRAS-WT (Figure 3A) and KRAS-G12V mutant complexes with RAF1 (Figure 3B) highlighted long-range dynamic changes induced by oncogenic mutation in the P-loop. In particular, structural projection of representative conformations from MD trajectories illustrated subtle and yet noticeably reduced thermal fluctuations especially in the switch II region of the G12V mutant as compared to the KRAS-WT. Our results also suggest that G12V mutation can affect the allosteric communication between switch I and switch II. The stabilized active conformation of switch II reduces its ability to interact dynamically with switch I, leading to a more rigid and less cooperative interaction. This mutation seems to indirectly reduce the flexibility of switch I, as the stabilized switch II region limits the conformational changes that can propagate to switch I. As a result, the reduced flexibility and altered interactions between switch I and switch II may limit the ability of KRAS to adopt alternative conformations, which may affect its interaction with other regulatory proteins.

**Figure 3.**
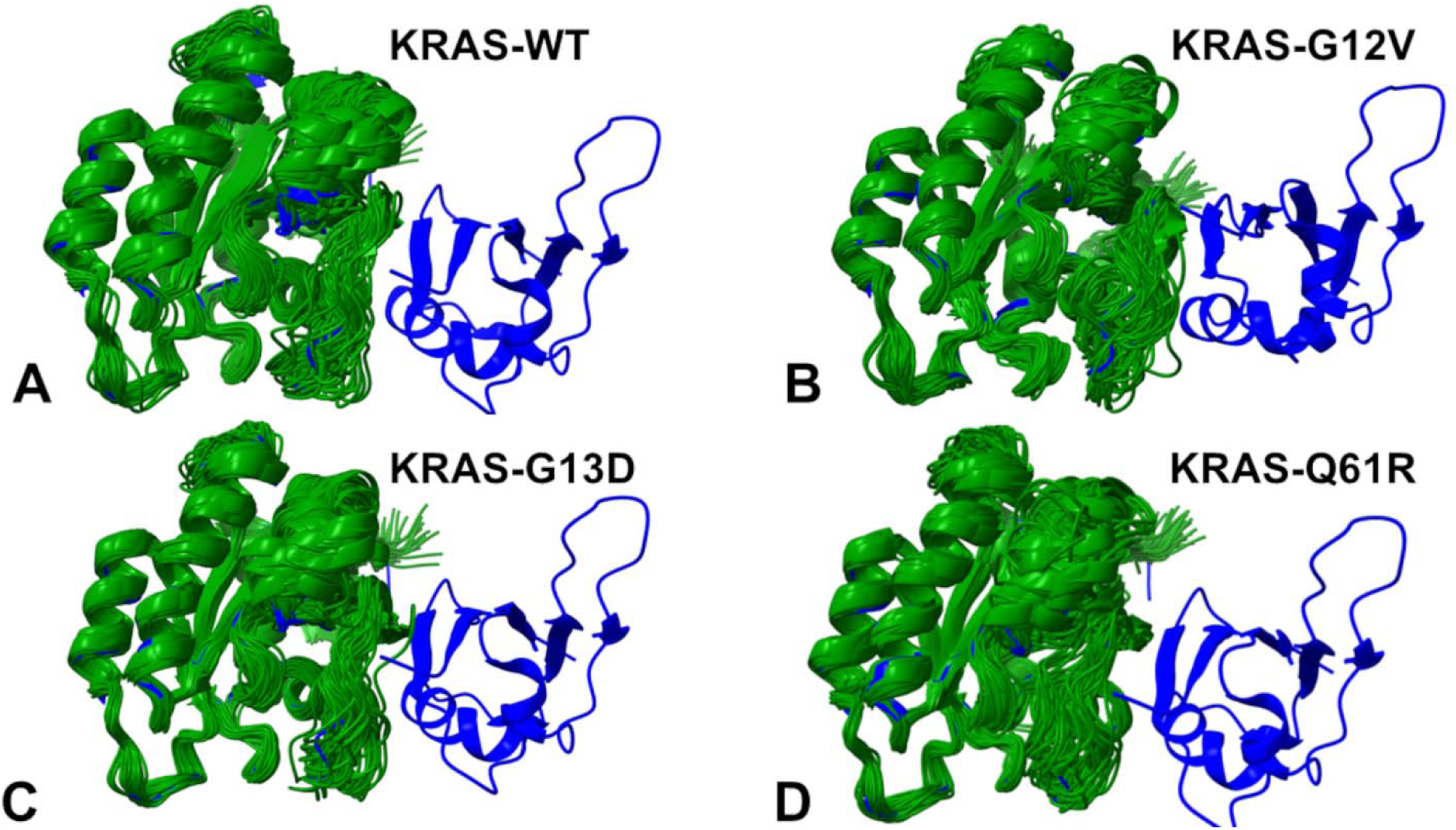
Structural mapping of the MD snapshots taken each 100 ns three independent 2 µs MD simulations of the crystal structure of wild-type KRAS (GMPPNP-bound) in complex with the RBD of RAF1/CRAF (pdb id 6VJJ); (A), KRAS-G12V mutant in the complex with the RBD of RAF1/CRAF (pdb id 6VJJ); (B), crystal structure of KRAS-G13D (GMPPNP-bound) in complex with the RBD and CRD of RAF1/CRAF (pdb id 6XGV) (C) and the crystal structure of KRAS-Q61R (GMPPNP-bound) in complex with RBD and CRD of RAF1/CRAF (pdb id 6XGU) (D). The KRAS conformational samples are shown in dark green ribbons and overlayed over the crystal structure of the KRAS-WT complex with RBD RAF1 (shown in blue ribbons).

G13D is another critical residue that can be mutated in cancer. We employed the crystal structure of KRAS-G13D (GMPPNP-bound) in complex with the RBD and CRD of RAF1/CRAF (pdb id 6XGV) and also the KRAS-G13D modeled mutant based on the crystal structure of the KRAS-WT bound to RAF1 (pdb id 6VJJ). MD simulations showed that G13D mutation in KRAS moderately reduces fluctuations in the switch I region while moderately increasing flexibility of the switch II region as seen in the KRAS-WT (Figure 2C). Our results suggested that G13D mutation may moderately reduce switch I fluctuations due to stabilization of the P-loop through electrostatic interactions with K117 residue. These results are consistent with structural studies revealing that the switch regions of KRAS show minor conformational changes in the structures of oncogenic mutants G13D complexed with RBDCRD when compared with WT KRAS-RBDCRD structure.^20^ Interestingly, the RMSF profiles (Figure 2C) and structural analysis of the conformational ensemble for KRAS-G13D complex (Figure 3C) indicated that mobility of the switch II region may be restored back to the KRAS-WT level.

Although these dynamic changes in the KRAS-G13D are fairly moderate and rather subtle, a detailed inspection of the conformational ensembles pointed to stabilization of the open active form in which the stability of the switch I is coupled to moderate fluctuations of switch II (Figure 3C). The observed functional differences among various oncogenic mutants are likely rooted in subtle conformational and dynamic differences. Similar to the NMR studies^90^, we observed that D13 side chain can sample several rotameric conformations and interact with positively charged K117 in the allosteric lobe ( residues 87-166) of KRAS, thus stabilizing the open state of switch I required for productive binding with the effector protein. NMR experiments revealed synchronized conformational dynamics in the active (GMPPNP-bound) KRAS-G13D and exchange between two conformational states in solution.^90^ NMR detected states of KRAS-G13D ( state I and state II) corresponded respectively to the closed and open position of the dynamic switch I relative to the bound nucleotide.^90^ It is worth noting that in the known conformational space of KRAS, the differences between state 1 and state 2 conformations in the crystal structures are localized to the switch I region (residues 30–38), corresponding to the closed/open forms and the switch II region (residues 60–76), showing significant conformational variability among various crystal structures, while the rest of the KRAS backbone is highly similar across all known structures.^90–94^

Our results showed that in the complex with RAF1, KRAS-G13D adopts stable open conformation of switch I while allowing functionally relevant variability in switch II (Figure 2C, 3C). The increased flexibility of switch II facilitates its interaction with the RBD of RAF1, as the region can more easily adopt the active conformation required for binding. It may also enhance the dynamic allosteric communication between switch I and switch II. The increased flexibility of switch II allows it to interact more dynamically with switch I, promoting a more cooperative interaction. Our results are consistent with other computational studies^31^ that indicated the ability of oncogenic mutations on the P-loop G12C, G12V and G13D can increase the flexibility of switch II region. The Q61R mutation stabilizes the open conformation of SWI by inducing conformational changes in the switch II region, which are propagated to switch I (Figure 2D). The RMSF profiles for the KRAS-Q61R mutant showed the increased variability of switch II while the thermal fluctuations of the open configuration of switch I are suppressed (Figure 2D). The stabilized open conformation of switch I assures the fidelity of its interaction with the RBD of RAF1, as key residues (e.g., Y32, D33, E37) are more consistently exposed for binding. Together, the results seem to suggest that all three mutations stabilize or enhance the active conformation of switch I, leading to increased binding to the RBD of RAF1 (Figure 2). We found that Q61R mutation can increase the flexibility of the switch II region of KRAS (Figure 2D).

Hence, MD simulations of Q61R-mutated KRAS revealed the increased flexibility in the switch II region and the reduced flexibility in the switch I region. While similar pattern was found for G13D mutant, the variability of switch II becomes more pronounced for Q61R mutant, thus potentially further promoting active open form of switch I (Figure 3D). The recent NMR relaxation dispersion experiments ^95^ showed that Q61R leads to a complete shift towards the active state and behaves differently from the oncogenic mutations as Q61R side-chain points towards switch I and anchors to switch I residues P34, T35 and I36 (Figure 3D). Our simulations explain this by observing of a transient hydrogen bond between the R61 side chain and the T35 backbone carbonyl oxygen that impose extra stabilization force on the open conformation of switch I without impeding variability of switch II region (Figure 2D, 3D). The increased mobility of switch II in the Q61R mutant can reduce the dynamic coupling with switch I leading to a more stable conformation of switch I which serves as the primary interface for KRAS-RAF1 interactions. The stable switch I conformation enhances the interaction with the RBD of RAF1, which may have functional implications for prolonged signaling and oncogenic transformation. Our results corroborate with these experiments suggesting that balance between rigidification of switch I and flexibility of switch II can promote shift towards the active state compatible with RAF1 binding. These findings are also consistent with the earlier studies^13,96^ showing that a hyperdynamic switch-II induces a more stable interaction between switch-I of KRAS and RAF1.

The RAF1 RBD is a structurally dynamic domain composed of a four-stranded antiparallel β-sheet (residues 56-63, 66-73, 96-101,125-130) and a single α-helix (residues 74-89). The core of the RBD composed of a β-sheet and α-helix structure remains highly stable throughout the interaction with KRAS (Figure 4).

**Figure 4.**
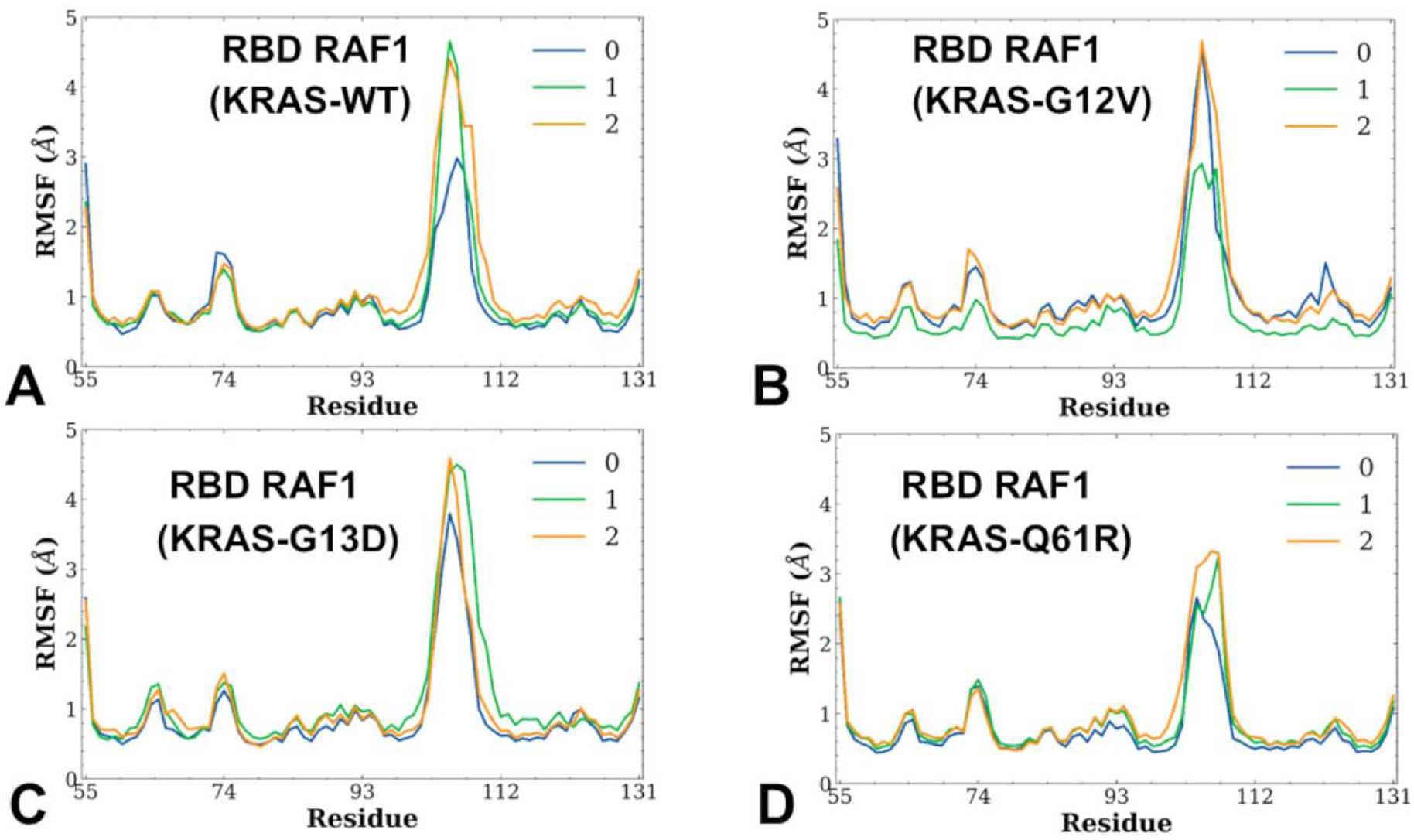
Conformational dynamics profiles for the RBD of RAF1 obtained from three independent MD simulations of the KRAS complexes with RAF1. The RMSF profiles for the RBD residues obtained from three independent 2 µs MD simulations of the crystal structure of wild-type KRAS (GMPPNP-bound) in complex with the RBD of RAF1/CRAF (pdb id 6VJJ); (A), KRAS-G12V mutant in the complex with the RBD of RAF1/CRAF (pdb id 6VJJ); (B), crystal structure of KRAS-G13D (GMPPNP-bound) in complex with the RBD and CRD of RAF1/CRAF (pdb id 6XGV) (C) and the crystal structure of KRAS-Q61R (GMPPNP-bound) in complex with RBD and CRD of RAF1/CRAF (pdb id 6XGU) (D).

The flexible loops (residues 51-55, 84-88,) allow the RBD to adapt to the conformational changes in KRAS during binding. The moderate flexibility of the RBD loops, combined with the stability of its core structure, allows the RBD to effectively interact with the flexible switch regions of KRAS (Figure 4). This dynamic yet stable interaction is essential for the proper activation of RAF1/CRAF. The ability of the RBD to maintain its structural integrity while accommodating the conformational changes in KRAS ensures that the interaction can be both specific and robust, enabling precise regulation of cellular signaling.^2^

To quantify dynamic couplings and correlations between motions of the KRAS regions we performed the dynamic cross correlation (DCC) residue analysis and reported the DCC maps for the KRAS-RAF1 complexes (Figure 5). By comparing the DCC maps of KRAS-WT (wild-type) and its oncogenic mutants (G12V, G13D, and Q61R), we can gain insights into how these mutations alter the dynamic couplings and correlations between motions of different KRAS regions. We observed KRAS-WT generally positive correlations between the P-loop (residues 10-17), Switch I (residues 25-40) and Switch II regions (residue 60-76) reflecting their functional coupling during nucleotide binding and hydrolysis (Figure 5A). The α-helices ( α-helix 1: 15–24; α-helix 2: 67–73; α-helix 3: 87–104; α-helix 4: 127–136; α-helix 5: 148–166) and β-sheets ( β-strand 1: 3–9; β-strand 2: 38–44; β-strand 5: 109–115; β-strand 6: 139–143) showed moderate correlations with the switch regions, while anti-correlated motions are observed between β-sheets (β-strand 3: 51–57; β-strand 4: 77–84) and the allosteric lobe ( residues 87-166) of KRAS (Figure 5A). The G12V mutation introduces a bulky hydrophobic residue in the P-loop. The reduced flexibility of switch II in the KRAS-G12V results in stronger positive correlations between residues within switch II as the region becomes more rigid. The stabilization of switch II reduced its correlations with other regions, such as the P-loop and switch I as well as suppresses negative correlations between switch II and β-sheets (β-strand 3: 51–57; β-strand 4: 77–84) as the rigid switch II becomes less responsive to motions in these regions (Figure 5B).

**Figure 5.**
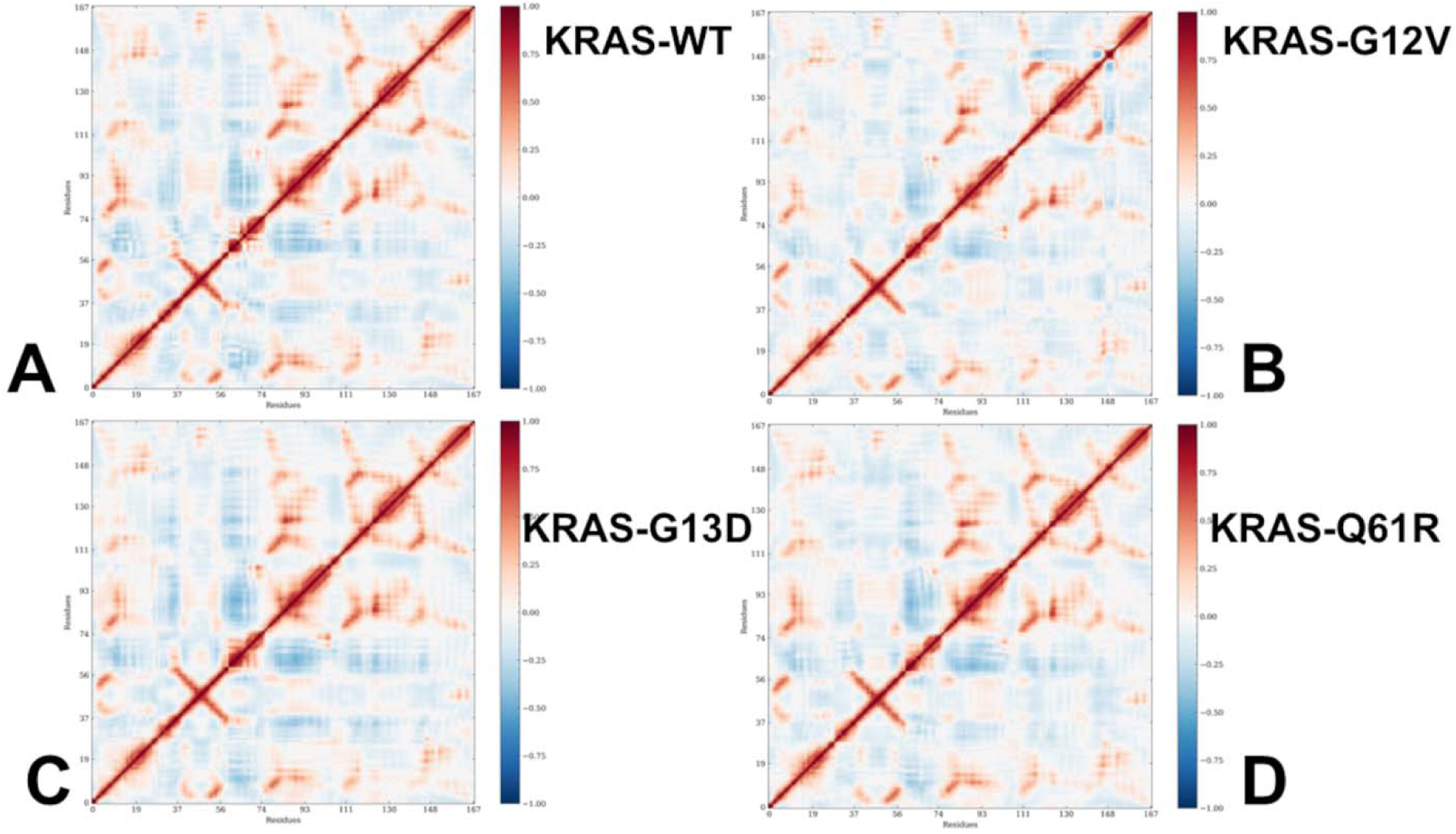
The DCC maps for the KRAS residues in the KRAS-WT complex with the RBD of RAF1/CRAF (pdb id 6VJJ); (A), KRAS-G12V mutant complex with the RBD of RAF1/CRAF (pdb id 6VJJ); (B), KRAS-G13D complex with the RBD and CRD of RAF1/CRAF (pdb id 6XGV) (C) and KRAS-Q61R complex with RBD and CRD of RAF1/CRAF (pdb id 6XGU) (D).

DCC measures the correlation between the motions of pairs of residues over the course of MD simulation. Positive correlations indicate that residues move in the same direction, while negative correlations indicate anti-correlated motions.

The DCC map for KRAS-G13D featured the increased positive correlations within switch II residues (Figure 5C) and a stronger pattern of anti-correlated couplings between dynamic switch II and other regions including allosteric lobe of KRAS. The increased flexibility of switch II facilitates dynamic allosteric communication with switch I and promotes a more cooperative interaction. Although DCC maps for G13D (Figure 5C) and Q61R KRAS mutants (Figure 5D) look quite similar, there are some subtle dynamic differences. In particular, the increased mobility of switch II in the Q61R mutant can reduce positive correlations between switch I and switch II, reflecting the weakened dynamic coupling between these regions. In addition, a more stable switch I conformation is constrained in the open form through transient hydrogen bond between the R61 side chain and the T35 backbone carbonyl oxygen. As a result, we observed weakened anti-correlated dynamic couplings between switch I and other regions (Figure 5D). The rigidification of switch I and a reduced dynamic coupling with switch I is a consequence of a shift towards the active state, which is compatible with RAF1 binding and prolonged signaling.

### Dimensionality Reduction Analysis and Markov State Modeling of KRAS Conformational Ensembles Reveals Principal Dynamic Consequences of Oncogenic Mutations

While the analysis of MD simulations and functional movements provided important insights into the underlying conformational landscape the high dimensionality of the data sets produced by simulations often hinders salient dynamic signatures associated with the mechanisms of allosteric transitions. Here, to facilitate the conformational landscape analysis we employed a dimensionality reduction method to project the results of MD simulations into low dimensional space.^97–100^ Given a time-series of molecular coordinates provided by the MD trajectories, tICA aims to reduce the dimensionality of the trajectories and to identify hidden key structural changes.

We leveraged conformational ensembles obtained from MD simulations for MSM analysis to estimate probabilities for protein transition among different macrostates. To build MSM, dimensionality reduction process needs to retain information about how proteins transition among macrostates. We analyzed the relaxation timescales in MSM, also referred to as implied timescales. The relaxation timescale can be interpreted as the time needed for a system to change its state. To cluster different conformations into metastable states, *k*-means clustering method was used to build clusters with the mean RMSD within cluster smaller than 1 Å. In the t-ICA reduced 2D space, k-means clustering method was applied to partition the 2D data. The top timescales are shown (Figure 6A). The trend of implied relaxation time scale revealed that the estimated time scale converged after approximately 10 ns, which was chosen as the lag time in the construction of MSM.

**Figure 6.**
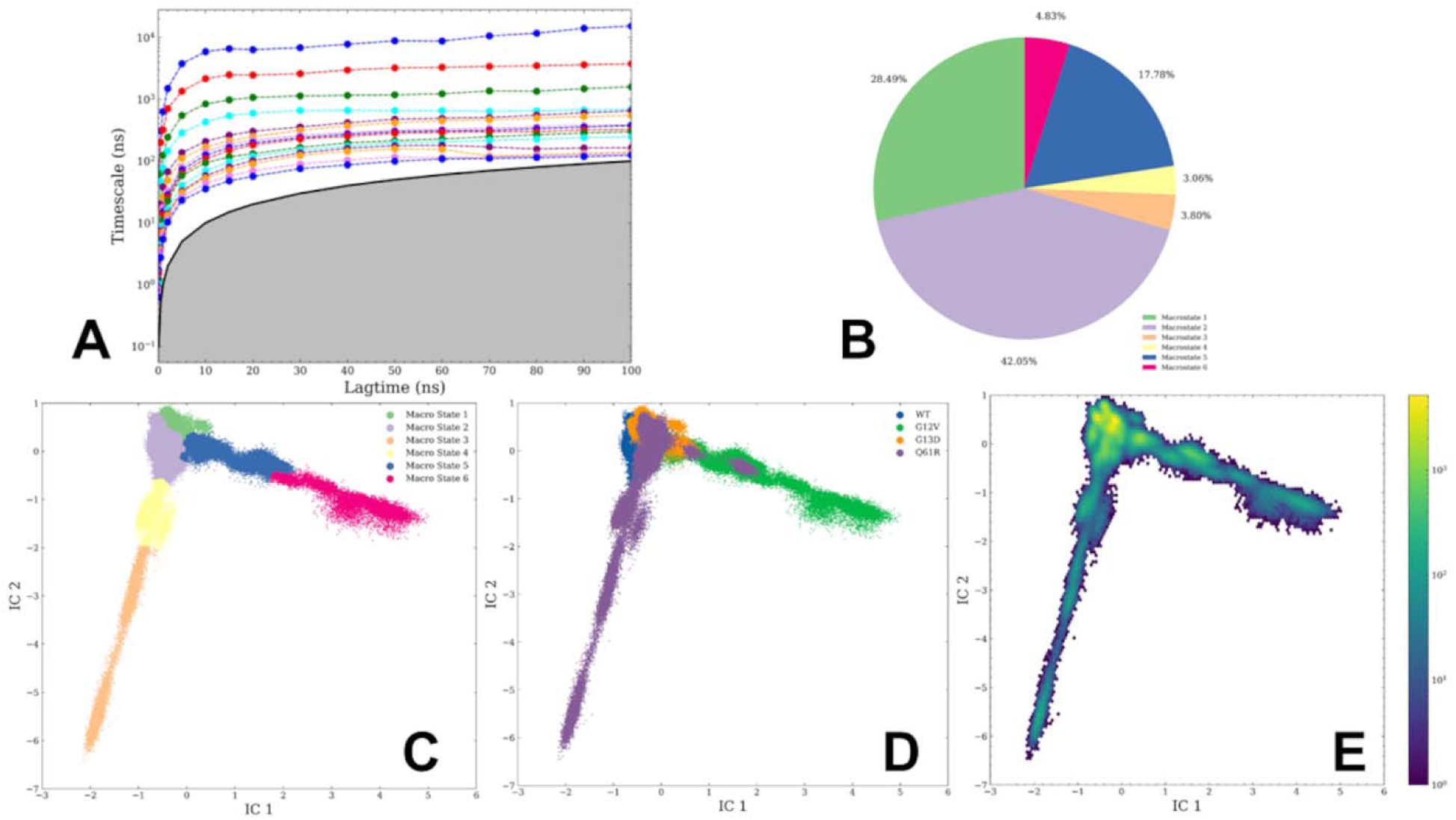
MSM analysis of the conformational landscapes for the KRAS-RAF1 complexes. (A) The estimated relaxation timescale is based on the transition probabilities among different microstates using different lag times. The top 15 timescales are shown. Each line represents the implied timescale as a function of lag time, and it is based on the corresponding eigenvalue. The estimated time scale converged after ∼10 ns, which was chosen as the lag time in the construction of MSM. The grey area depicts a “linear” region where lag time (x axis) equals the relaxation timescale (y axis) but note that y axis is shown in log scale. The implied timescales cross under the grey region correspond to processes which are faster than the lag time. (B) The pie diagram of the composition and occupancy of the macrostates. (C) The low-dimensional density representations with 2 components (t-IC) are shown for macrostates 1-6. (D) The low-dimensional density representation with 2 components (t-IC). The distribution densities are colored based on the system and are shown for KRAS-WT complex, KRAS-G12V, KRAS-G13D and KRAS-Q61R complexes. (E) The distribution density of all simulations is visualized using a hexagon bin plot. Each hexagon represents a small region, and the color bar shows the frequency of this region.

The density plot revealed that several macrostates (microstate 1, 2, and 5) are dominant and correspond to 28.49 %, 42.05 %, and 17.78% of the entire population of states obtained from MD simulations of all systems (Figure 6B,C). Interestingly, the low-dimensional projection of conformations showed that the macrostates 1 and 5 occupy different “legs” of the distribution and may reflect different conformational signatures associated with semi-open and open KRAS conformations. The low-dimensional map of the KRAS-WT and KRAS mutant complexes highlighted a more restricted conformational space afforded by KRAS-G12V and KRAS-G13D mutants that appear to lock switch I and II in the open active conformation, while larger and more diverse conformational landscape is induced by Q61R mutation that exerts activation through different dynamic mechanisms and redistribution of the conformational landscape (Figure 6D).

The hexagon binning was employed to represent the distribution density of all MD simulations, with each hexagon representing a small region, and the color bar showing the frequency of this region (Figure 6E). Since differences in the conformational space of KRAS are localized to the switch I region corresponding to the closed, semi-open and active open forms ^90–94^ we infer that the distinct signatures of major macrostates may indeed reflect this principal pattern of the KRAS dynamics (Figure 6). After the partition of the MSM macrostates was determined the stationary distribution and transition probabilities were calculated based on the constructed MSM (Figure 7). The transition probabilities are determined among different macrostates for all systems. The high percentage of self-conserved probability shows the stability of macrostates (Figure 7). MSM identifies a dominant macrostate 2 (93%) corresponding to the fully active conformation of KRAS-WT, where both switch I and switch II are ordered and engaged with the RBD of RAF1 (Figure 7A). In addition, the KRAS-WT populates macrostates 1,4 and 5. For KRAS-G12V, the distribution of macrostates is similar, with dominant microstate 2 (77%) and macrostates 1,5 and 6 that occupy only one “leg” of the distribution (Figure 7B). These macrostates reflect semi-open transient conformations that KRAS-WT samples during the transition between the active and inactive states in which switch I fluctuates between almost-open and full open conformations (Figure 7B) MSM analysis of KRAS-G13D demonstrated even a narrower range of transitions between the dominant microstates 2 and 1 that have the high percentage of self-conserved probability (72% for microstate 2 and 85% for microstate 1) reflecting the stability of these macrostates (Figure 7C).

**Figure 7.**
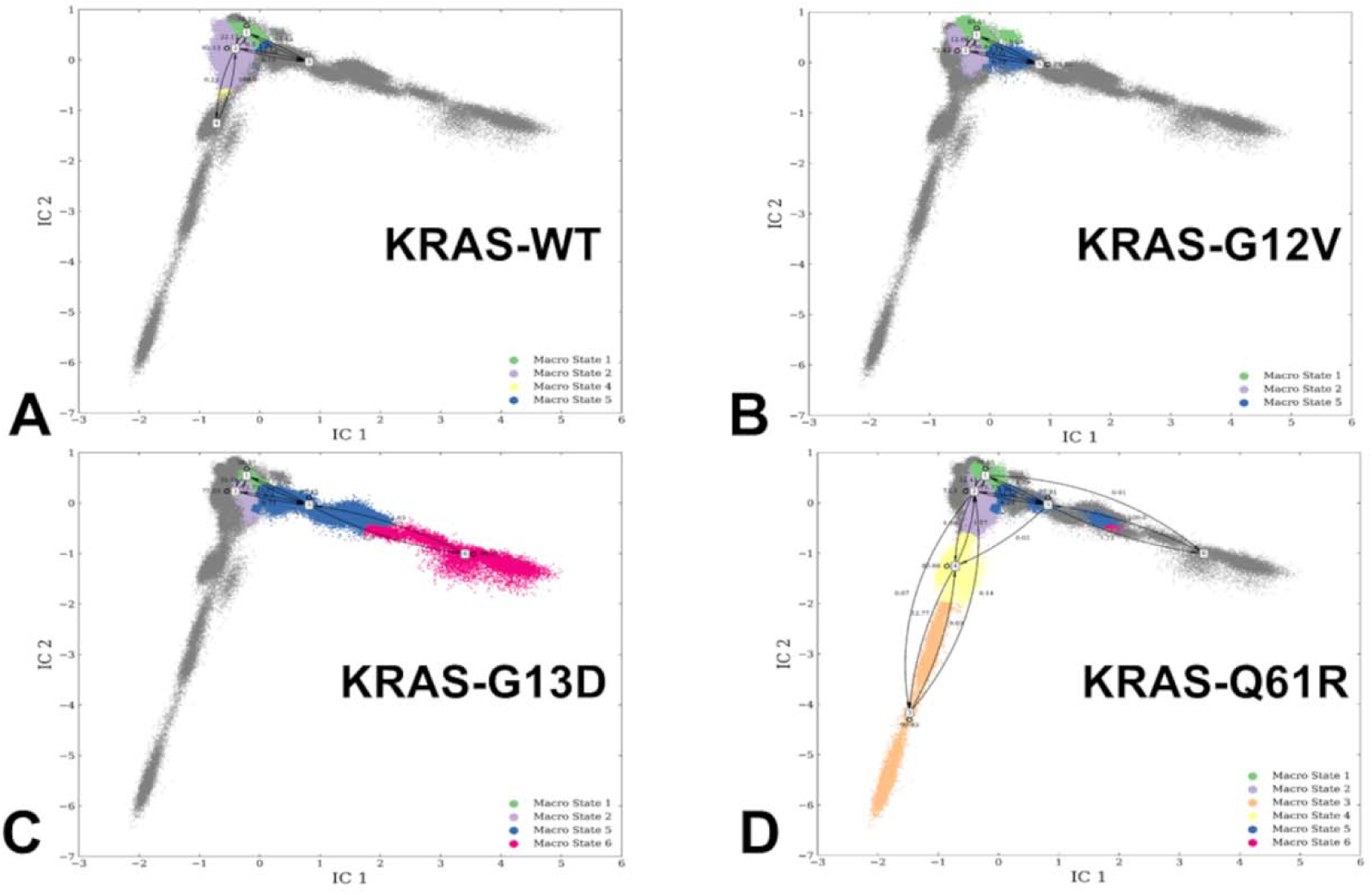
MSM analysis of the conformational landscape for the KRAS-WT complex, KRAS-G12V, KRAS-G13D and KRAS-Q61R complexes. The transition probability maps among different macrostates with the 5 ns lag time for KRAS-WT complex (A), KRAS-G12V (B), KRAS-G13D (C) and KRAS-Q61R complexes (D). The high percentage of self-conserved probability shows the stability of macrostates.

Both macrostates featured stable open conformation of switch I and moderate variability in the active switch II region. The vastly different distribution is obtained for KRAS-Q61R complex with RAF1 revealing expansion of macrostates and reflecting the increased conformational plasticity of KRAS (Figure 7D). Indeed, we observed that in addition to main macrostates 1 and 2 there is significant population of other macrostates (3,4,5, and 6) (Figure 7D). The transition probabilities are especially high between macrostates 1 and “right leg” of the distribution (macrostates 5,6). Notably, macrostates 5 and 6 are also found for KRAS-G13D (Figure 7C,D). However, unique to Q61R, the distribution also featured macrostates 3 and 4 with smaller and yet appreciable transition probabilities to microstate 2 (Figure 7D). The macrostates 3,4 correspond to semi-open intermediate conformations that are not significantly populated in the wild-type or other mutant forms of KRAS. These intermediate states represent semi-open configurations of switch I and transiently stable conformations of switch II. Hence, the Q61R mutation partly alters the energy landscape of KRAS, lowering the energy barriers between macrostates and allowing for more frequent transitions between them which results in a more complex and dynamic conformational landscape (Figure 7D). The expansion of macrostates induced by the Q61R mutation may have important structural and functional implications for KRAS signaling. The expanded number of macrostates reflects the increased dynamic behavior of KRAS, which may facilitate its interaction with a broader range of effectors. The stabilization of intermediate states and the increased flexibility of switch II may also prolong the interaction between KRAS and RAF1, leading to prolonged signaling and enhancing effector binding. NMR studies of KRAS mutants reveal the dynamic nature of the switch regions and the existence of multiple conformational states,^90^ consistent with the MSM results.

To conclude, MSM analysis revealed that KRAS mutants may exhibit subtle and yet distinct conformational dynamics signatures compared to KRAS-WT, with stabilization of the active state and altered flexibility of the switch regions. These dynamic signatures of KRAS mutants may be relevant in modulating and enhancing the binding affinity of KRAS for downstream effectors RAF1. Overall, the MSM analysis provides useful insights into conformational dynamics and potential relevance for functional impact, highlighting the importance of targeting the dynamic properties of KRAS for therapeutic intervention.

### Mutational Scanning of KRAS-RAF1 Protein Binding and Stability Reveals Conserved Energy Hotspots and Effects of Oncogenic Mutants

Using conformational ensembles obtained from MD simulations, we performed systematic mutational scanning of the KRAS residues in the KRAS-RAF1 complexes (Figure 8). In silico mutational scanning was done by averaging the binding free energy changes over the equilibrium ensembles and allows for predictions of the mutation-induced changes of the binding interactions and the stability of the complex. To provide a systematic comparison, we constructed mutational heatmaps for the KRAS interface residues. We used BeatMusic^69–71^ and PRODIGY contact predictor^101,102^ to identify binding interface residues. Residues are considered part of the interface if they are within a defined cutoff distance (typically 5 Å) from atoms in the binding partner. Our definition of binding interface residues corresponds to the definition of the contact residues in the experimental studies.^29^ The mutational heatmap of the KRAS-RAF1 complex revealed several important binding affinity hotspots that correspond to I36, Y40, R41 and Y64 residues (Figure 8A). The largest destabilization changes are associated with mutations of Y40 including Y40D ( ΔΔG = 2.67 kcal/mol), Y40S ( ΔΔG = 2.49 kcal/mol), Y49E ( ΔΔG = 2.45 kcal/mol) and Y40K ( ΔΔG = 2.3 kcal/mol) (Figure 8A). Consistent with the free energy experiments^29^, this simplified model established the importance of the aromatic side chain in Y40 KRAS position which makes a cation–π interaction with RAF1 R89 residue. The results of mutational scanning recapitulated subtle experimental data showing that E37D mutation is not tolerated while E37 replacements by Y, F and H can be tolerated indicating that the salt bridge to RAF1 R67 can be substituted by interactions involving an aromatic side chain (Figure 8A).

**Figure 8.**
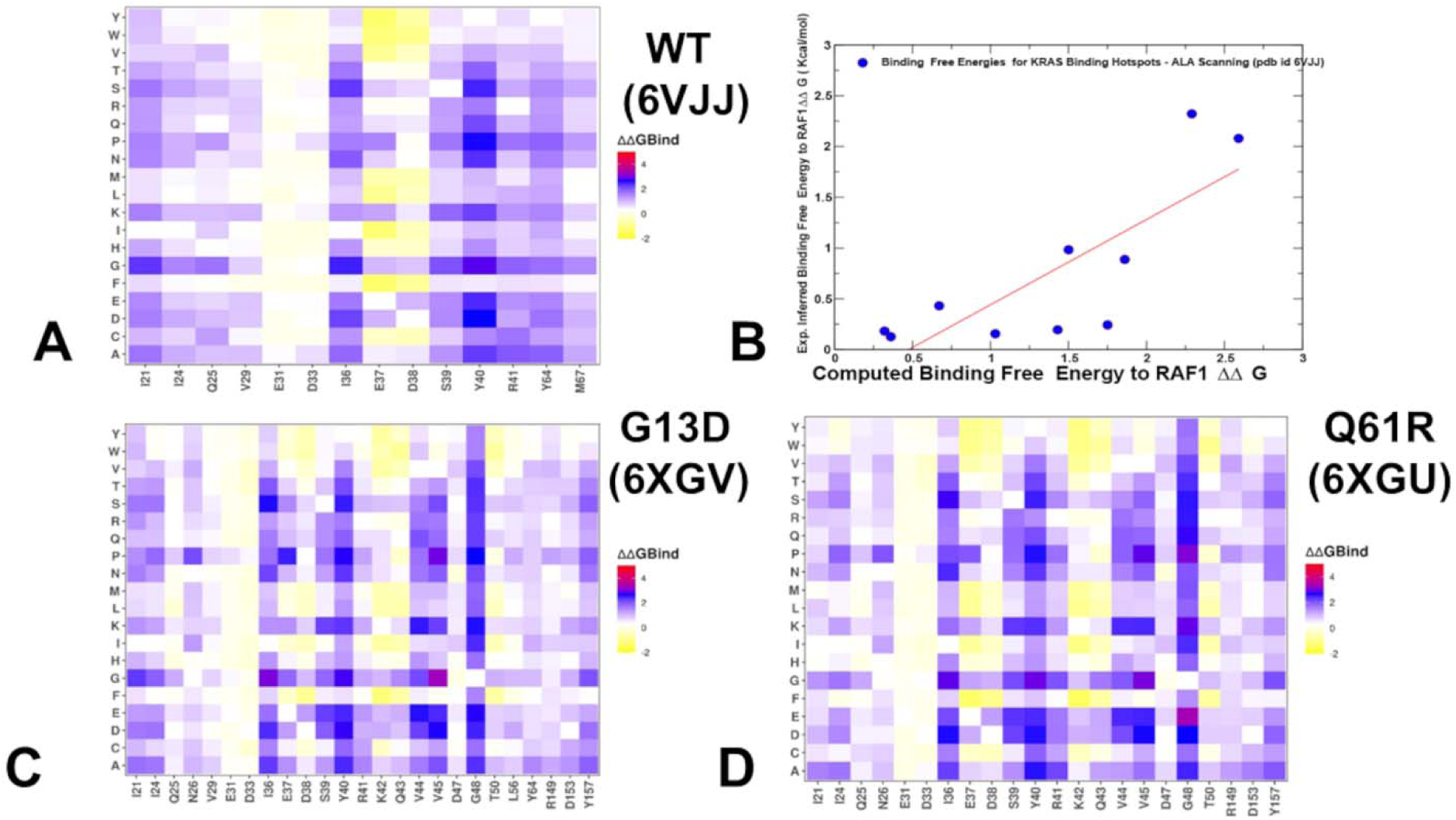
Ensemble-based dynamic mutational profiling of the KRAS intermolecular interfaces in the KRAS-RAF1 complexes. The mutational scanning heatmaps are shown for the interfacial KRAS residues in the KRAS-WT complex (A). A correlation between experimentally inferred binding energies^29^ and predicted binding free energy changes for a group of experimentally established KRAS binding hotspots that included residues I21, I24, E31, D33, I36, E37, D38, S39, Y40 and R41 (B). The mutational scanning heatmaps are shown for the interfacial KRAS residues in the KRAS-G13D (C) and KRAS-Q61R complexes (D). The heatmaps show the computed binding free energy changes for 20 single mutations of the interfacial positions. The standard errors of the mean for the binding free energy changes using randomly selected 1,000 conformational samples from the MD trajectories were 0.11-0.18 kcal/mol,

Consistent with the functional mutational scanning data^29^, the mutational heatmap also showed that I21, I36 and S39 are important binding hotpot that makes multiple hydrophobic contacts with RAF1 and where polar mutations are highly detrimental (Figure 8A). As was found experimentally, mutations at other interface residues that contact RAF1 are better tolerated, particularly mutations at D33 with only charge-reversing mutations to R and K and mutation to P mildly inhibiting binding (Figure 8A). Similarly, mutations at the other two charged site E31 at the edge of the interface have essentially neutral effect on the binding free energy (Figure 8A). The only position where mutational scanning disagreed with the experiments is D38 where according to our data, only reversing charge mutations cause detrimental effects while other mutations can be tolerated (Figure 8A). In contrast, the experimental data showed that D38 cannot be changed to any other amino acid without detrimental effects on binding affinity.^29^

We also directly compared the experimental and predicted binding free energy changes for a group of experimentally established KRAS binding hotspots hotspots that included residues I21, I24, E31, D33, I36, E37, D38, S39, Y40 and R41 (Figure 8B). This analysis revealed a statistically significant correlation between computed and experimental binding free energy changes, supporting the results and interpretations of computational studies performed in this study.

Using MD ensembles obtained from simulations of and KRAS G13D complex with RBD-CRD (pdb id 6XGV) and KRAS-Q61R complex with RBD-CRD of RAF1/CRAF (pdb if 6XGU) we conducted mutational scanning of the KRAS interface residues (Figure 8C,D). The KRAS binding interface is expanded as the interfacial residues include now interactions with both RBD and CRD proteins. Importantly, we found that the effects of G13D and Q61R mutations on the binding affinity and interface residues of RBD RAF1 is relatively moderate. Indeed, the KRAS binding hotspots I36, Y40, R41 are preserved but notably mutations at E37 positions become less tolerant while modifications at R41 are slightly less detrimental albeit still unfavorable (Figure 9C,D). G13D mutation introduces a negatively charged aspartate residue, which impairs GTP hydrolysis and locks KRAS the GTP-bound (active) state, leading to constitutive activation. By locking KRAS in an active state, G13D enhances the accessibility of the switch I and II regions for RAF1 binding. While the G13D mutation does not directly alter the E37 and D38 residues in the switch I region, it appeared to enhance their interactions with RAF1 by stabilizing the active conformation of KRAS (Figure 8C). E37 forms a salt bridge with R67 of RAF1 and electrostatic interactions with R59 which is critical for stabilizing the KRAS-RAF1 interaction. Also located in the switch I region, D38 interacts with R67 and R89 of RAF1 through electrostatic interactions. Although the mutational changes in E37 positions of the G13D KRAS complex are not dramatic, they are appreciable and may result from G13D-induced stabilization of the active conformation which enhances the accessibility of E37 and D38 for interaction with RAF1 (Figure 8C).

**Figure 9.**
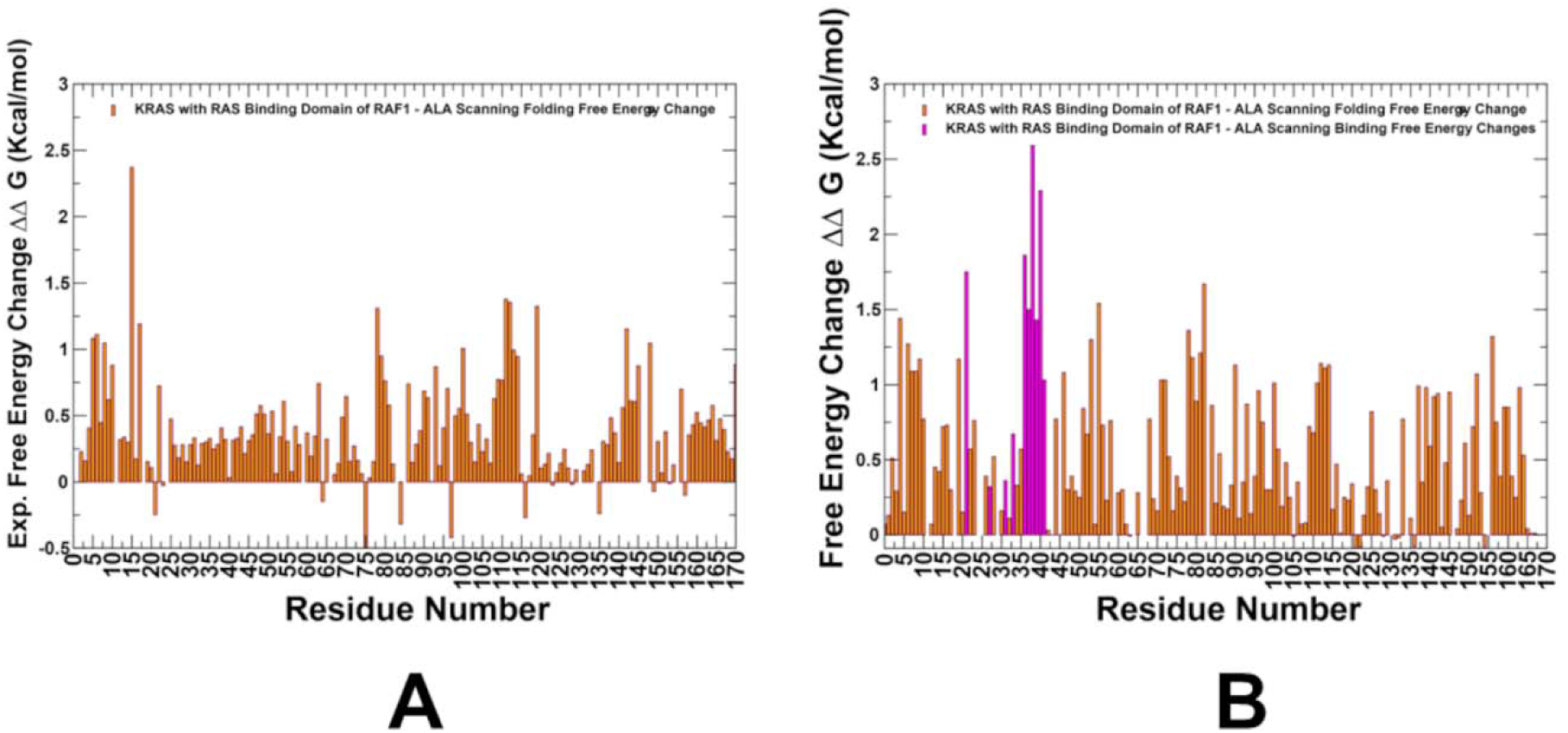
A comparison of the experimentally determined binding free energy changes upon alanine mutational scanning of the KRAS residues^29^ (A) and the computed folding and binding free energy changes upon alanine mutational scanning of the KRAS residues (B). The folding free energy changes are shown in orange-colored filled bars in (A) and (B). The computationally predicted binding KRAS-RAF1 binding free energy changes associated with alanine substitutions are shown in magenta-colored filled bars on panel (B).

Mutational heatmap for the KRAS-Q61R mutant complex displayed generally very similar patterns as G13D with some stronger effect on binding of I36 and Y40 and modestly weaker effect on E37 and D38 (Figure 8D). Structural studies^20^ and functional experiments^29^ showed that Q61 mutations stabilize the active conformation of KRAS, enhancing the binding contributions of key hotspots like I36, Y40, and R41. Indeed, consistent with these experimental studies, the mutational heatmap for KRS-Q61R complex showed that key binding hotspots are I36, S39, Y40 and R41 for RBD of RAF1 and additionally V45 and G48 positions that are binding affinity hotspots for interactions with CRD of RAF1 (Figure 8D). In contrast to the KRAS-RBD interface consisting of polar and charged interactions, the KRAS-CRD interface contains no salt bridges and large hydrophobic interface. KRAS interacts with CRD mainly via its residues present in the inter-switch region (R41, K42, Q43, V44, V45, I46, D47, and G48), and in the C-terminal helix α5 (R149, D153, and Y157). Mutational heatmaps showed that V44, V45 and G48 are critical hotspots of binding with CRD (Figure 8C,D). V44 interacts with hydrophobic residues in the RBD and CRD of RAF1, such as L154 and V155. In particular, mutations at this position (e.g., V44A or V44D) could disrupt hydrophobic packing and showed to be highly detrimental for binding (Figure 8B,C). The mutational heatmaps pointed also to KRAS residues R149, D153, and Y157 that are interfacing specifically with the CRD RAF1 (Figure 9C,D). These residues at the C-terminal helix of KRAS and are peripherally positioned near the CRD of RAF1. However, this region contributes to the overall stability of the KRAS-RAF1 interface, even though it is not part of the switch I or switch II regions. R149 is a positively charged residue that can form electrostatic interactions with acidic residues in the CRD, while D153 is engaged in electrostatic interactions H175, R179 of the CRD of RAF1. Mutations at this position (D153A or charge reversal D153K, D153R) disrupt these interactions, reducing binding affinity (Figure 8C,D). Finally, Y157 mutations have stronger detrimental effects on binding affinity ad this aromatic residue participates in hydrophobic interactions and hydrogen bond interactions with E174 of the CRD of RAF1 (Figure 8C,D). Overall, the interactions formed by these KRAS residues with CRD are weaker and less significant for binding than those provided by V44, V45 and G48 sites. As a result, mutations in in R149, D153, and Y157 are more tolerant to mutations as suggested by the computed heatmaps. These results are consistent with the DMS data^29^ indicating that besides the KRAS residues present in the inter-switch region, especially residues K42 to V45, play an important role in RAS-mediated RAF activation, most likely through KRAS-CRD interactions (Figure 8C,D).

In addition, we performed structure-based mutational scanning of KRAS binding interface in other complexes with other binding partners, including KPIK3CG (pdb id 1HE8) (Supporting Information, Figure S3A), RALGDS (pdb id 1LFD) (Supporting Information, Figure S3B), SOS1 (pdb id 1NVW) (Supporting Information, Figure S3C), DARPin K27 (pdb id 5O2S) (Supporting Information, Figure S3D), and DARPin K55 (pdb id 5O2T) (Supporting Information, Figure S3E) and compared the results with the experimental data. We first considered how mutations in the binding interfaces alter binding to four additional interaction partners. These proteins bind KRAS through an overlapping set of contacts and shared contacts which is particularly pronounced for the three effector proteins, RAF1 (Figure 8A), PIK3CG (Supporting Information, Figure S2A) and RALGDS (Supporting Information, Figure S2B). By comparing mutational effects, we found that some residues particularly I21, I36, Y40 and Y64 are critically important for binding to all three proteins (Supporting Information, Figure S2). For example, many mutations of hydrophobic residues I36 and Y40 strongly inhibit binding to all three effectors. In general, mutational scanning against all partners showed relative tolerance of KRAS residues E31 and D33 and also similar patterns for mutations of more vulnerable positions E37 and D38 (Supporting Information, Figure S2). The residues E37 and D38 are shared binding hotspots in KRAS, critical for interactions with RAF1, PIK3CG, and RALGDS. Mutations at these residues (e.g., E37D, E37A, D38A, D38K) significantly reduce binding affinity, while certain substitutions (e.g., E37Y/F/H, D38E) may be tolerated depending on the binding partner. Many substitutions at E37 were experimentally shown to inhibit binding to RAF1 and RALGDS but increase binding to PIK3CG. Mutations of S39 have stronger effect on binding to PIK3CG and RAF1 but not to RALGDS. Hence, although mutational scanning can accurately and robustly predict major binding affinity hotspots and the effects of most detrimental mutations, the resolution of simplified knowledge-based energy function may be often insufficient to capture more subtle and sensitive specificity changes particularly in some polar interface residues.

To get deeper understanding of protein stability effects, we examined the folding free energy changes for KRAS residues that are not involved in the binding interface to identify protein stability hotspots and compare the results with the experiments. According to the experiments, the major peaks of stability changes in the alanine scanning are associated with KRAS residues K5, L6, V8, G10, G15, K16, S17, F78, L79, V81, I100, M111, V112, D119, T142, T148 (Figure 9A).^29^ The P-loop also known as the G-loop is a highly conserved region in GTPases like KRAS. It plays a critical role in binding and hydrolyzing GTP, which is essential for KRAS function. The residues G10, G15, S17, and S19 in the P-loop are particularly important for the folding and stability of KRAS. The P-loop must be highly flexible to accommodate GTP binding and hydrolysis. Glycine residues (G10, G15) are essential for this flexibility, as they allow the loop to adopt the necessary conformations. The computational alanine scanning of all KRAS residues showed that most detrimental effects on protein stability occur upon mutations of residues F82, K5, L6, V8, G10, L79, V81, V112, L113, V114 (Figure 9B). Hence, a simplified energy model is capable of reproducing important stability changes and identifying most of the stability hotspots in agreement with the experimental data. We identified the key stability centers in the P-loop as well as F78, L79 and V81 positions that are part of the hydrophobic core of KRAS, which is essential for maintaining the protein’s overall stability and folding. The KRAS core defined by relative accessible surface area < 0.25 includes α-helix 1: 15–24; α-helix 2: 67–73; α-helix 3: 87–104; α-helix 4: 127–136; α-helix 5: 148–166; β-strand 1: 3–9; β-strand 2: 38–44; β-strand 3: 51–57; β-strand 4: 77–84; β-strand 5: 109–115; β-strand 6: 139–143.^29^ β-strand 4 (residues 77-84) is located in the central β-sheet of KRAS structure which is critical for maintaining the protein’s structural integrity.^20,29^ Our results showed that disruption of F78, L79 and V81 residues from this region through mutation to alanine destabilizes the hydrophobic core, leading to misfolding and reduced stability of KRAS (Figure 9B).

Overall, the results suggested that mutations detrimental to folding are enriched in the hydrophobic core of the protein. The study highlights that mutations detrimental to folding are enriched in the hydrophobic core and P-loop of KRAS. These regions are critical for maintaining the protein’s stability and function, and mutations in these regions are likely to have significant functional consequences. The computational model successfully reproduced the experimental stability hotspots, demonstrating its ability to identify key residues critical for KRAS stability. This agreement validates the use of the proposed computational alanine scanning as a tool for predicting protein stability effects in the KRAS complexes.

### MM-GBSA Analysis of the Binding Energetics Provides Quantitative Characterization of Thermodynamic Drivers of Protein Interactions

Using the conformational equilibrium ensembles obtained MD simulations of the KRAS-RAF1 WT and KRAS mutant complexes we computed the binding free energies for these complexes using the MM-GBSA method.^72–75^ MM-GBSA is also employed to (a) decompose the free energy contributions to the binding free energy of a protein–protein complex on per-residue basis; (b) evaluate the role of hydrophobic and electrostatic interactions as thermodynamic drivers of KRAS-RAF1 binding; (c) identify the binding energy hotspots and compare the rigorous MM-GBSA predictions of binding energetics with the results of mutational scanning of KRAS residues in the KRAS complexes.

We started with the MM-GBSA analysis of the KRAS-WT complex with RBD of RAF1 (Figure 10A-C). The energy decomposition showed that the binding affinity hotspots correspond to switch I residues S39 (ΔG = −6.77 kcal/mol), Y40 (ΔG = −5.8 kcal/mol). D38 (ΔG = −5.45 kcal/mol), D33 (ΔG = −4.98 kcal/mol), R67 (ΔG = −4.77 kcal/mol) and R41 (ΔG = −4.1 kcal/mol) (Figure 10A). The analysis of the van der Waals contributions pointed to a critical role of this binding energy component for hotspot Y40 (ΔG_VDW_ = −4.74 kcal/mol), R41 (ΔG_VDW_ = −3.41 kcal/mol), E37 (ΔG_VDW_ = −3.33 kcal/mol) and S39 (ΔG_VDW_ = −2.34 kcal/mol) (Figure 10B).

**Figure 10.**
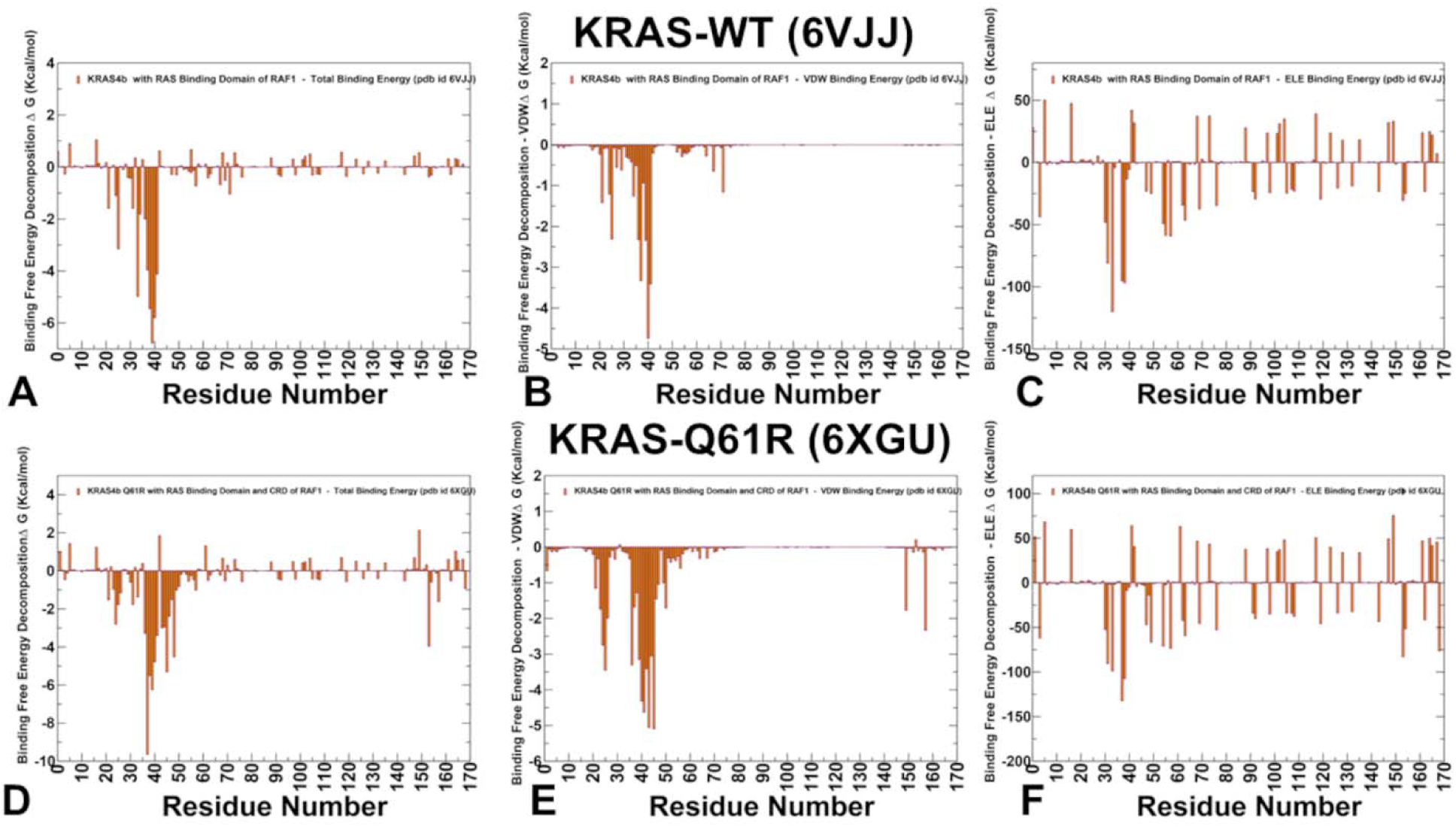
The residue-based decomposition of the total binding MM-GBSA energies for the KRAS residues in the KRAS-WT complex with RBD RAF1 (A) and KRAS-Q61 mutant (D). The residue-based decomposition of the van der Waals contribution to the total MM-GBSA binding energy for the KRAS residues in the KRAS-WT complex with RBD RAF1 (B) and KRAS-Q61R mutant (E). The residue-based decomposition of the electrostatic contribution to the total MM-GBSA binding energy for the KRAS residues in the KRAS-WT complex with RBD RAF1 (C) and KRAS-Q61R mutant (F). The MM-GBSA contributions are evaluated using 1,000 samples from the equilibrium MD simulations of respective KRAS-RAF1 complexes. It is assumed that the entropy contributions for binding are similar and are not considered in the analysis.

Of particular significance is the contribution of electrostatic interactions as some major binding hotspots of KRAS correspond to the charged residues E37, D38, D33, E168 and D57 (Figure 10C). Overall. MM-GBSA calculations strongly suggested that both van der Waals, and electrostatic interactions work synergistically to yield the most favorable binding energies for Y40, E37, D38, S39 and R41 hotspots (Figure 10A). These results are in excellent agreement with the sophisticated biochemical and DMS studies^29^ unequivocally showing that the interface residues that are most important for RAF1 binding include a mixture of charged (E37 and D38) and hydrophobic (I36 and Y40) residues. Y40 forms hydrophobic interactions and a cation-π interaction with R89 of RAF1. E37 forms a salt bridge with R67 of RAF1 and this electrostatic interaction is critical for stabilizing the switch I region and enhancing binding affinity (Figure 10A-C). For comparison, we also highlight the energy decomposition results for KRAS-Q61R mutant complex with RB-CRD of RAF1 (Figure 10D-F). The key binding affinity hotspots with RBD of RAF1 in this complex are E37 (ΔG = −9.64 kcal/mol), S39 (ΔG = −6.25 kcal/mol), D38 (ΔG = −6.25 kcal/mol), and Y40 (ΔG = −4.78 kcal/mol) (Figure 10D). In addition, the results revealed top binding hotspots for KRAS binding with CRD of RAF1 which are V45 (ΔG = - 5.31 kcal/mol) and G48 (ΔG = −4.52 kcal/mol) (Figure 10D). The most favorable van der Waals interactions are formed by R41, Y40, I36 and S39 with RBD of RAF1, while V45 binding to CRD is driven primarily by the strongest van der Waals contacts (Figure 10E). Similarly to KRAS-WT, KRAS-Q61R binding to RBD is driven by strong electrostatic interactions of switch I residues E37, D38 and D33 as well as another strong interacting center D57 from β-strand 3 ( 51–57). Hence, consistent with the experiments, our results similarly established that the central β-sheet plays a key role in modulating KRAS activity and particularly mediating allosteric binding with RAF1 (Figure 10F).

Overall, a comparison of MM-GBSA energies for KRAS-WT and KRAS-Q61R mutant complexes indicated that a conserved pool of major KRAS binding centers from switch I particularly E37, D38 and Y40 are dominant binding affinity hotspots shared among studied KRAS mutants. Importantly, we found very similar energetics patterns and the same binding hotspots for KRAS-G13D complex with RBD-CRD of RAF1 (Supporting Information, Figure S4).

To further leverage the MM-GBSA energetic analysis, we performed systematic mutational scanning of the key interfacial residues E31, D33, E37, D38, S39, Y40 and R41 (Figure 11). The results clearly showed significant destabilizing changes upon all mutations of key binding hotspot residues D33, E37, D38 and Y40 (Figure 11B,C,D,F). In particular, mutational scanning underscored the role of electrostatic interactions as the dominant contribution of D33 to the KRAS-RAF1 binding interface (Figure 11B). D33 forms a salt bridge with K84 of RAF1, which is a strong electrostatic interaction and electrostatic contacts with N71. As a result, the electrostatic interactions serve as the primary thermodynamic driver of D33 binding hotspot, and the disruption of this interactions particularly by charge reverse mutations D33K, D33R reduces binding affinity significantly (Figure 11B) which is precisely what was found experimentally.^29^

**Figure 11.**
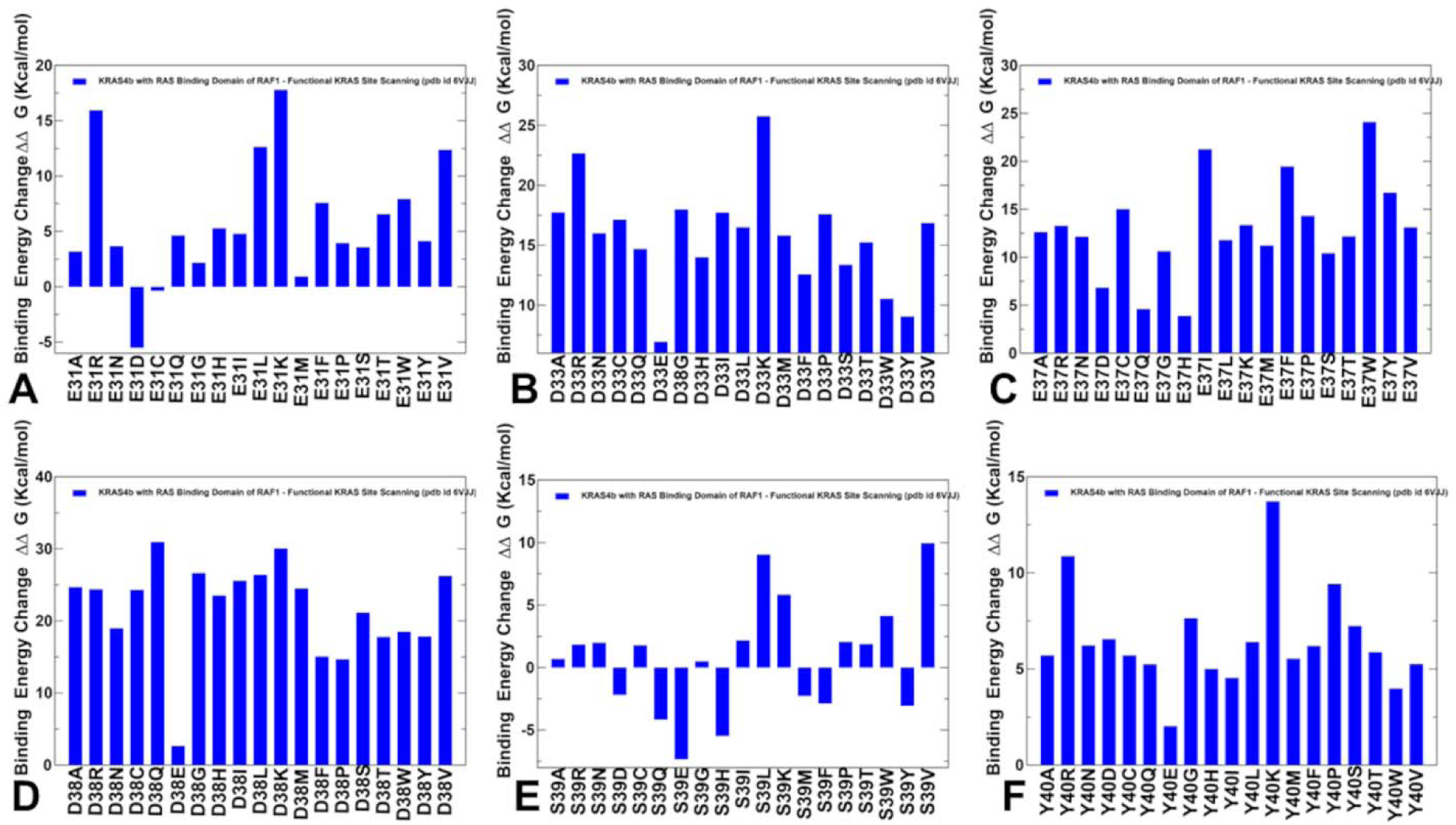
MM-GBSA mutational scanning of the KRAS binding interface residues with RBD RAF1. Mutational scanning results are shown for KRAS binding hotspots E31 (A), D33 (B), E37 (C), D38 (D), S39 (E) and Y40 (F). The binding free energy changes are shown in magenta-colored filled bars. The positive binding free energy values ΔΔG correspond to destabilizing changes and negative binding free energy changes are associated with stabilizing changes.

Similarly, MM-GBSA based mutational scanning of E37 hotspot reflected the role of strong salt bridges formed by E37 with R59 and R67 residues on RBD RAF1 (Figure 11C). The interactions of E37 with R67 and R59 stabilize switch I region, which is critical for high-affinity binding to RAF1.

This stabilization ensures that the switch I region adopts the proper conformation for interaction with RAF1. In addition, E37 makes favorable hydrophobic contacts with T68 and V69 which amounts to significant van der Waals contribution (ΔG_VDW_ = −3.33 kcal/mol). The largest destabilizing energy changes upon mutations were observed for two major hotspots D38 (Figure 11D) and Y40 (Figure 11F). D38 makes strong electrostatic interactions with R67 and R89 and also favorable van der Waals contacts with A85, T68 and V69 of the RAF1. As a result, such mutations as D38Q, D38K and D38V result in large destabilization changes (Figure 11D). This is consistent with experiments showing that D38 cannot be changed to any other amino acid without detrimental effects on binding affinity.^29^ Similarly, Y40 forms highly favorable cation– π interaction with RAF1 R89 residue as well as favorable contacts with V88 and Q66 of RAF1. For Y40 hotspot, mutations Y40K and Y40R also produced large destabilizing changes (Figure 11F). Interestingly, the binding energy changes upon mutations are considerably smaller for E31 and S39, with some mutations at S39 position such as S39Q and S39E resulting in a more favorable binding with RAF1 (Figure 11A,E). Hence, MM-GBSA analysis and mutational profiling showed that the hotspots can be grouped in the top tier (E37, D38, Y40) and secondary tier (E31 and S39). These observations are fully consistent with the functional experiments showing that Y40 and D38 are indeed the most dominant binding hotspots in which not a single mutation can be tolerated for binding affinity with RAF1.^29^

To summarize the MM-GBSA computations and analysis, the MM-GBSA analysis highlights the synergistic role of electrostatic and hydrophobic interactions in stabilizing the KRAS-RAF1 complex. Residues lY40 and E37 contribute to binding through both types of interactions, underscoring the importance of a balanced interplay between these forces. The results also highlighted the critical role of switch I residues D33, E37, D38, and Y40, playing a critical role in binding to RAF1. The MM-GBSA predictions are in excellent agreement with experimental data,^29^ underscoring the reliability of MM-GBSA for identifying binding energy hotspots and understanding the thermodynamic drivers of protein-protein interactions. The MM-GBSA based mutational scanning results provide insights into the effects of specific mutations on KRAS-RAF1 binding, particularly charge-reversal mutations (e.g., D33K, D33R, D38K, Y40K) that significantly destabilize the complex, highlighting the importance of electrostatic interactions. Importantly, the MM-GBSA results are also consistent with mutational scanning and heatmaps produced by simplified energy models. Both approaches revealed conserved binding mechanism across KRAS variants. Indeed, the identification of conserved binding hotspots (E37, D38, Y40) across KRAS-WT, KRAS-Q61R, and KRAS-G13D suggests a shared binding mechanism. This conservation highlights the importance of these residues in mediating KRAS-RAF1 interactions, regardless of the specific mutation. The results overall align well with experimental data and provide a detailed understanding of the thermodynamic drivers of KRAS-RAF1 interactions.

### Probing Allostery in the KRAS-RAF1 Complexes Using Dynamic Network Analysis : Allosteric Landscape of KRAS Binding and Hotspots of Allosteric Communications

We used the ensemble-based network centrality analysis and the network-based mutational profiling of allosteric residue propensities that are computed using the topological network parameter SPC to characterize global network of allosteric communications. Through ensemble-based averaging over mutation-induced changes in the SPC metric, we identify KRAS positions in which mutations on average cause network changes. Allosteric hotspots are identified as residues in which mutations incur significant perturbations of the global residue interaction network that disrupt the network connectivity and cause a significant impairment of global network communications and compromise signaling. By performing in silico version of “deep” mutational scanning to measure the allosteric effects in the KRAS-RAF1 we examine the variant-induced network changes.

Using a graph-based network model of residue interactions we first computed the ensemble-averaged distributions of the residue-based SPC metric (Figure 12A). SPC is a measure of the influence of a node in a network based on the number of shortest paths that pass through it. Nodes with high betweenness centrality function as bridges or bottlenecks in the network, facilitating communication between different parts of the network. The distributions revealed several critical clusters of residues that are important for mediating allosteric communications, particularly concentrated in regions (residues 6-8, 15-25, 35-43, 53-60, 75-80, 95-100, 115-120) (Figure 12A). These important for allosteric communication sites include residue from P-loop, switch-I, switch-II, β-strand 4: 77–84 α-helix 3: 87–104 and β-strand 5: 109–115. The dominant peaks correspond to residues L6, I55, G15, K16, S17, E37, D38, S39, D57, M72, F78, H95, Y96, R97 (Figure 12A).

**Figure 12.**
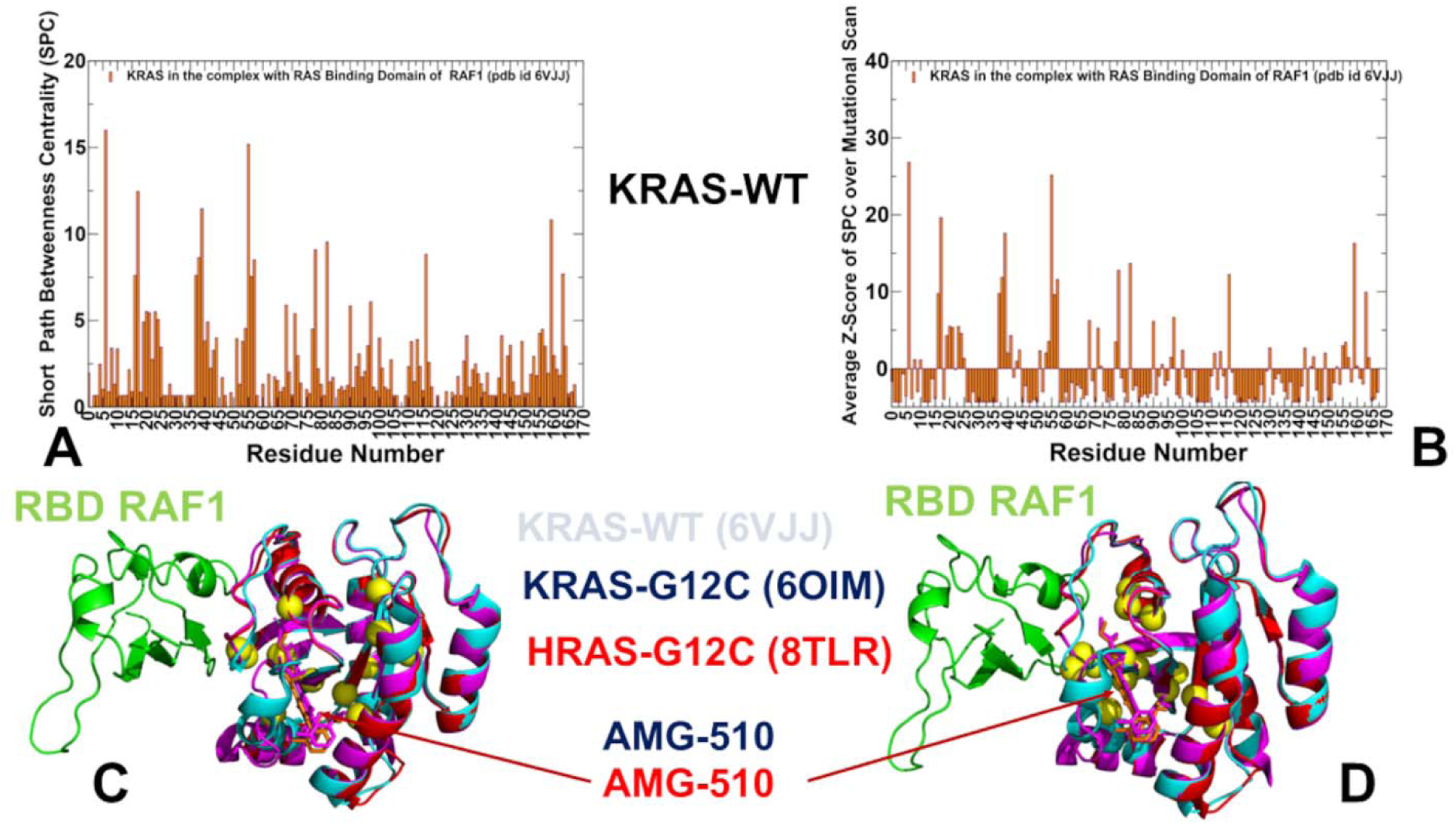
(A) The ensemble-averaged SPC centrality for the KRAS residues in the KRAS-WT complex with RAF1 ( in orange filled bars). (B) The SPC peaks (G15, K16, S17, E37, D38, S39, I55, D57, M72, F78, H95, Y96, and R97 residues) defining potential allosteric centers are shown in yellow spheres and mapped onto the crystal structure of the KRAS-RAF1 complex (pdb id 6VJJ). The KRAS-WT is in cyan ribbons. The KRAS-RAF1 complex is overlayed with the structures of human KRAS-G12C bound with allosteric inhibitor AMG510 ( blue ribbons, AMG50 is in blue sticks) and human HRAS-G12C bound to AMG510 (red ribbons, AMG510 is in red sticks). (C) The Z-score of SPC centrality for KRAS residues averaged over mutational scan for each residue ( in orange filled bars). (D) Structural mapping of the predicted Z-score peaks and assigned allosteric hotspots with KRAS positions S17, S39, L159, F82, F78, N116, D38 D57, R97 and R68 shown in yellow spheres. Similar to panel (B), the structures of KRAS-RAF1 (cyan ribbons), human KRAS-G12C complex (blue ribbons) and HRAS-G12C complex (red ribbons) are shown along with the bound allosteric inhibitor AMG510.

By mapping these sites onto the crystal structure of the KRAS-RAF1 complex, we found that the predicted allosteric centers are lined up to form major allosteric communication routes connecting KRAS core with the KRAS-RAF1 binding interface (Figure 12B). One group of residues is located in a small groove that is known to be occupied by allosteric inhibitors,^16^ while another cluster consists of key binding interface residues from switch I (E37, D38, S39, Y40). Strikingly, the predicted allosteric cluster that is distant from the RAF1 binding interface consists of KRAS residues ((H95, Y96, R97) forming cryptic allosteric pocket that is targeted by known allosteric inhibitor sotorasib (AMG 510) which is the first KRAS(G12C) inhibitor to reach clinical testing in humans.^16^ Indeed, structural studies showed that the isopropyl-methylpyridine substituent of AMG 510 occupied the His95 groove engaged in a network of primarily van der Waals contacts extending from the backbone of helix 2 (H95, Y96) to the flexible switch II loop (Supporting Information, Figure S5). The recent illuminating study from Shokat lab used structural and pharmacological analysis of members of the Ras, Rho, and Rab family of GTPases to demonstrated that this cryptic allosteric pocket in the switch II region of KRAS can be present in other GTPases beyond KRAS indicating that GTPases exhibit targetable allosteric binding pockets.^103^

We also examined the Z-score of SPC centrality for RBD residues averaged over mutational scan for the KRAS residues (Figure 12C). In this model, we characterize residues where mutations on average induce a significant increase in the SPC metric and therefore have a dramatic effect on the efficiency of long-range communications in the allosteric interaction network. This analysis enables identification of allosteric control points that could determine the efficient and robust long-range communications in the KRAS-RAF1 complexes. The profile further highlighted the major peaks in which mutational perturbations cause significant changes in allosteric communications. The emerging dominant peaks corresponded to KRAS positions L6, I55, S17, S39, L159, F82, F78, N116, D38 D57, R97 and R68 (Figure 12C). Structural map of Z-score peaks that define in this model potential allosteric centers reveals a dominant allosteric route connecting the central core of KRAS with the switch II region in the binding interface, while several residues including R68, Y96 and R97 are nestled in a more peripheral to the interface cryptic pocket targeted by allosteric drugs (Figure 12D). Notably, the network analysis did not explicitly compute the mutational effects in these allosteric centers on the binding affinity with RAF1 but rather on the efficiency of allosteric communications. Nonetheless, the striking revelation of the network analysis is that the majority of the predicted hotspots were also experimentally determined as allosteric binding hotspots.^29^ More specifically, among 18 experimentally identified allosteric binding centers that are distant from the binding interface with RAF1 and yet significantly affect binding affinity of KRAS are D57, G15, K16, S17 T35, F28, Y32, P34 positions^29^. Of particular significance are allosteric hotspots D57, G15, S17 that are away from the binding interface and yet induce significant destabilization in binding upon mutations. Our analysis identifies these sites as major hotspots of allosteric communication that affect interaction paths from remote parts of KRAS to the binding interface (Figure 12B,D).

These findings suggest that the efficiency of long-range communication between a given remote site and binding interface measured by profiling of the SPC metric is linked with the effect of this site on binding as mutations in the allosteric site can induce perturbations at the binding interface and thus impair binding with RAF1. Interestingly, one of the most striking findings of the experimental studies of KRAS allostery is the efficient propagation of allosteric effects across the central β-sheet of KRAS which acts as a hub for transmitting conformational changes linking distant functional sites.^29^ Consistent with these experimental findings, structural mapping of the predicted allosteric centers showed a significant population across the central β- sheet of KRAS. Indeed, the predicted hotspots L6 is from β-strand 1 (3–9), E37, D38, S39, Q43 belong to β-strand 2 (38–44), and I55, L56, D57 belong to β-strand 3 ( 51–57) (Figure 12B,D). Hence, in line with the experiments, our results similarly established that the central β-sheet plays a key role in modulating KRAS activity and particularly mediating allosteric binding with RAF1.

To summarize, the network-based modeling of allosteric interactions in KRAS-RAF1 complexes identified critical clusters of residues important for allosteric communication, including residues from the P-loop, switch I, switch II, β-strand 4 (77–84), α-helix 3 (87–104), and β-strand 5 (109–115). The predicted allosteric centers form major communication routes connecting the KRAS core with the KRAS-RAF1 binding interface. These routes connect the KRAS core with the switch I regions through a small groove occupied by allosteric inhibitors using the central β-sheet of KRAS for transmitting conformational changes and linking distant functional sites. Key residues in these routes include L6 (β-strand 1), E37, D38, S39 (switch I region), Q43 (β-strand 2), and I55, L56, D57 (β-strand 3).The predicted allosteric hotspots align with experimentally identified allosteric binding centers, such as D57, G15, K16, S17, T35, F28, Y32, and P34. These residues, although distant from the binding interface, significantly affect binding affinity with RAF1. The results also highlight the importance of allosteric communication in KRAS, where distant residues influence the binding interface with RAF1 through long-range interactions. This communication is mediated by a network of residues that function as bridges or bottlenecks in the allosteric network. The predicted allosteric hotspots, such as D57, G15, S17, and D38, represent potential therapeutic targets. Mutations in these residues disrupt allosteric communication and impair KRAS-RAF1 binding, highlighting their importance for maintaining KRAS function. Targeting these hotspots with small molecules could offer new strategies for inhibiting KRAS-driven signaling. A strong agreement between computational predictions and experimental data suggests that the network-based approach can be a robust predictor of allosteric binding hotspots.

## Discussion

The microsecond molecular dynamics (MD) simulations of KRAS-WT and its oncogenic mutants (G12V, G13D, and Q61R) in complex with RAF1 provide critical insights into the structural dynamics, conformational flexibility, and binding mechanisms of KRAS. These simulations reveal how oncogenic mutations alter the dynamic behavior of KRAS, The switch I region (residues 25-40) displays considerable flexibility, fluctuating between semi-open and fully open conformations. These fluctuations reflect the dynamic nature of switch I, which is critical for effector binding and signaling. The switch II region (residues 60-76) also exhibits flexibility, although it is more stable than switch I in the GTP-bound state. The Q61R mutation induces a more complex dynamic behavior, with increased flexibility in switch II and reduced flexibility in switch I. The simulations reveal that the Q61R mutation stabilizes the open conformation of switch I, ensuring that key residues (e.g., Y32, D33, E37) are consistently exposed for binding to RAF1. The increased flexibility of switch II in the Q61R mutant promotes dynamic allosteric communication between switches I and II enhancing RAF1 binding and signaling.

The MSM analysis highlights how oncogenic mutations (G12V, G13D, and Q61R) stabilize the active conformation of KRAS, enhancing its binding affinity for RAF1. This stabilization is achieved through reduced flexibility in switch I and II, which favors the open conformation required for effector binding. The stabilization of the active state contributes to the oncogenic behavior of KRAS mutants, leading to constitutive signaling and cell proliferation. The G12V mutation reduces the dynamic coupling between switch II and other regions, reflecting its rigidification. In contrast, the G13D and Q61R mutations enhance the flexibility of switch II, promoting cooperative interactions with switch I. These changes in allosteric communication highlight the importance of switch II as a mediator of KRAS dynamics and signaling. The expansion of macrostates induced by the Q61R mutation may have important structural and functional implications for KRAS signaling. The expanded number of macrostates reflects the increased dynamic behavior of KRAS, which may facilitate its interaction with a broader range of effectors.

Mutational scanning of KRAS-RAF1 complexes identified key binding affinity hotspots, including Y40, E37, D38, and D33, and together with the MM-GBSA analysis highlighted the synergistic role of electrostatic and hydrophobic interactions in stabilizing the KRAS-RAF1 complex. The key binding affinity hotspots E37, D38, Y40 contribute to binding through both types of interactions, underscoring the importance of a balanced interplay between these forces. These findings suggest that targeting these interactions could disrupt KRAS-RAF1 binding and inhibit oncogenic signaling. Similarly, the network-based analysis identified D57, G15, and S17 as allosteric hotspots, which have been experimentally shown to affect binding affinity despite being distant from the interface. The identification of binding affinity hotspots and allosteric communication routes provides a foundation for developing targeted therapies against KRAS-driven cancers. The electrostatic and hydrophobic interactions at the KRAS-RAF1 interface, particularly involving Y40, E37, and D38, are critical for binding and could be targeted by small molecules or peptides. These findings highlight the importance of integrating computational and experimental approaches to study KRAS dynamics and guide therapeutic intervention. The consistency between computational and experimental results underscores the importance of these methods for understanding KRAS dynamics and guiding therapeutic development.

While the computational studies presented in this work provide valuable insights into the allosteric regulatory landscapes of KRAS and its interactions with RAF1, there are several limitations that must be acknowledged. These limitations stem from the inherent challenges of modeling complex biomolecular systems and the approximations used in computational methods. These include timescale limitations of MD simulations, simplifications in MSM and network models, approximate energy functions in free energy calculations, Although the computational predictions align well with available experimental data there are still gaps in the experimental validation of certain predictions, particularly for allosteric hotspots and long-range communication pathways. For example, the predicted allosteric role of residues like D57 and G15 is supported by indirect evidence, but direct experimental validation (e.g., through mutagenesis or NMR relaxation experiments) is needed to confirm their functional importance. The timescales of MD simulations (nanoseconds to microseconds) are often much shorter than the timescales of experimental techniques (e.g., NMR relaxation, single-molecule FRET), which can measure dynamics on the millisecond to second timescale. This discrepancy can make it challenging to directly compare computational and experimental results. Despite these limitations, the findings of this study provide a strong foundation for understanding the allosteric regulation of KRAS and its interactions with RAF1.

## Conclusions

The results of this study provide important quantitative insights into principles of allosteric communication and allosteric binding in KRAS that are consistent with and can explain the experimental data. Microsecond MD simulations reveal that KRAS-WT and its oncogenic mutants (G12V, G13D, Q61R) exhibit unique conformational dynamics, particularly in the switch I and II regions. These mutations stabilize the active, open conformation of KRAS, enhancing RAF1 binding and promoting oncogenic signaling. Oncogenic mutations (G12V, G13D, Q61R) reduce flexibility in the switch regions, favoring the open, active state required for effector binding. This stabilization contributes to prolonged RAF1 interaction and constitutive signaling, driving oncogenic transformation. MM-GBSA and mutational scanning identify conserved binding hotspots (E37, D38, Y40) across KRAS variants and different binding partners, underscoring their critical role in mediating KRAS interactions with effector proteins. This conservation highlights the importance of these residues in mediating KRAS-RAF1 interactions, regardless of the specific mutation. The results overall align well with experimental data and provide a detailed understanding of the thermodynamic drivers of KRAS interactions. Network analysis identifies key allosteric communication routes, with residues like D57, G15, and S17 acting as critical hubs. These residues, though distant from the binding interface, significantly influence KRAS-RAF1 interactions, highlighting the importance of long-range allosteric regulation. The identification of binding affinity hotspots and allosteric communication routes provides a foundation for targeted therapeutic strategies. Disrupting key interactions (e.g., Y40, E37, D38) or targeting allosteric sites (e.g., D57, G15) could inhibit KRAS-driven signaling and offer new avenues for cancer treatment. The strong agreement between computational predictions and experimental data validates the use of MD simulations, MSM, and network analysis in studying KRAS dynamics. These methods offer robust tools for understanding KRAS allostery and guiding therapeutic development.

## Author Contributions

Conceptualization, G.V. and P.T.; methodology, S.X., G.V. P.T.; software, S.X., M.A., G.H., G.V and P.T.; validation, G.V.; formal analysis, G.V., M.A., G.H., S.X., and P.T. investigation, G.V. and P.T.; resources, G.V., M.A. and G.H.; data curation, G.V.; writing—original draft preparation, G.V.; writing—review and editing, S.X., G.V., M.A. and G.H.; visualization, G.V.; supervision, G.V.; project administration, P.T. and G.V.; funding acquisition, P.T. and G.V. All authors have read and agreed to the published version of the manuscript.

## Conflicts of Interest

The authors declare no conflict of interest. The funders had no role in the design of the study; in the collection, analyses, or interpretation of data; in the writing of the manuscript; or in the decision to publish the results.

## Funding

This research was supported by the National Institutes of Health under Award 1R01AI181600-01 and Subaward 6069-SC24-11 and the Kay Family Foundation Grant A20-0032 to G.V and National Institutes of Health under Award No. R15GM122013 to P.T.

## Data Availability Statement

Data is fully contained within the article and Supplementary Materials. Crystal structures were obtained and downloaded from the Protein Data Bank (http://www.rcsb.org). The rendering of protein structures was done with UCSF ChimeraX package (https://www.rbvi.ucsf.edu/chimerax/) and Pymol (https://pymol.org/2/). The software tools used in this study are freely available GitHub sites https://github.com/smu-tao-group/protein-VAE; https://github.com/smu-tao-group/PASSer2.0. All the data obtained in this work (including simulation trajectories, topology and parameter files, the software tools, and the in-house scripts are freely available at <u>ZENODO at</u> https://zenodo.org/records/13989093.

## Acknowledgments

G.V acknowledges support from Schmid College of Science and Technology at Chapman University for providing computing resources at the Keck Center for Science and Engineering.

## Notes

### Competing Interest Statement

The authors have declared no competing interest.

