## Supplemental Figures S1-S5 for "Accurate Characterization of the Allosteric Energy Landscapes, Binding Hotspots and Long-Range Communications for KRAS Complexes with Effector Proteins : Integrative Approach Using Microsecond Molecular Dynamics, Deep Mutational Scanning of Binding Energetics and Allosteric Network Modeling"

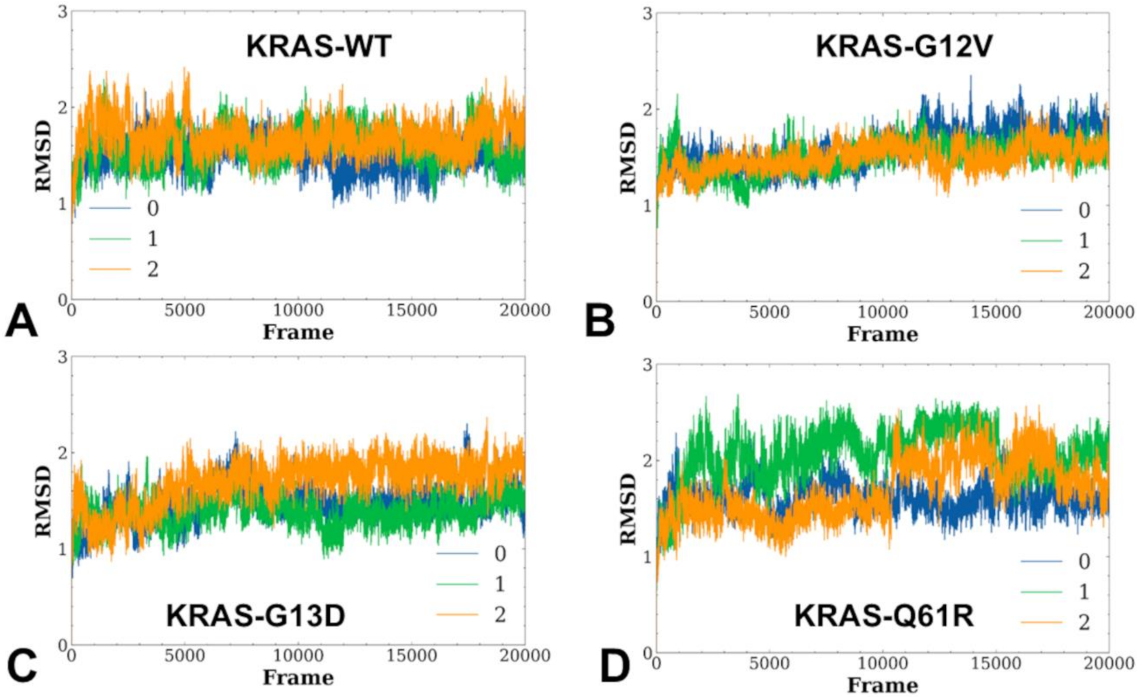


**Figure S1.** Conformational dynamics profiles of the KRAS protein residues obtained from three independent MD simulations of the KRAS complexes with RAF1. The RMSD profiles for the RBD residues obtained from three independent 2 μs MD simulations of the crystal structure of wild-type KRAS (GMPPNP-bound) in complex with the RBD of RAF1/CRAF (pdb id 6VJJ); (A), KRAS-G12V mutant in the complex with the RBD of RAF1/CRAF (pdb id 6VJJ); (B), crystal structure of KRAS-G13D (GMPPNP-bound) in complex with the RBD and CRD of RAF1/CRAF (pdb id 6XGV) (C) and the crystal structure of KRAS-Q61R (GMPPNP-bound) in complex with RBD and CRD of RAF1/CRAF (pdb id 6XGU) (D).

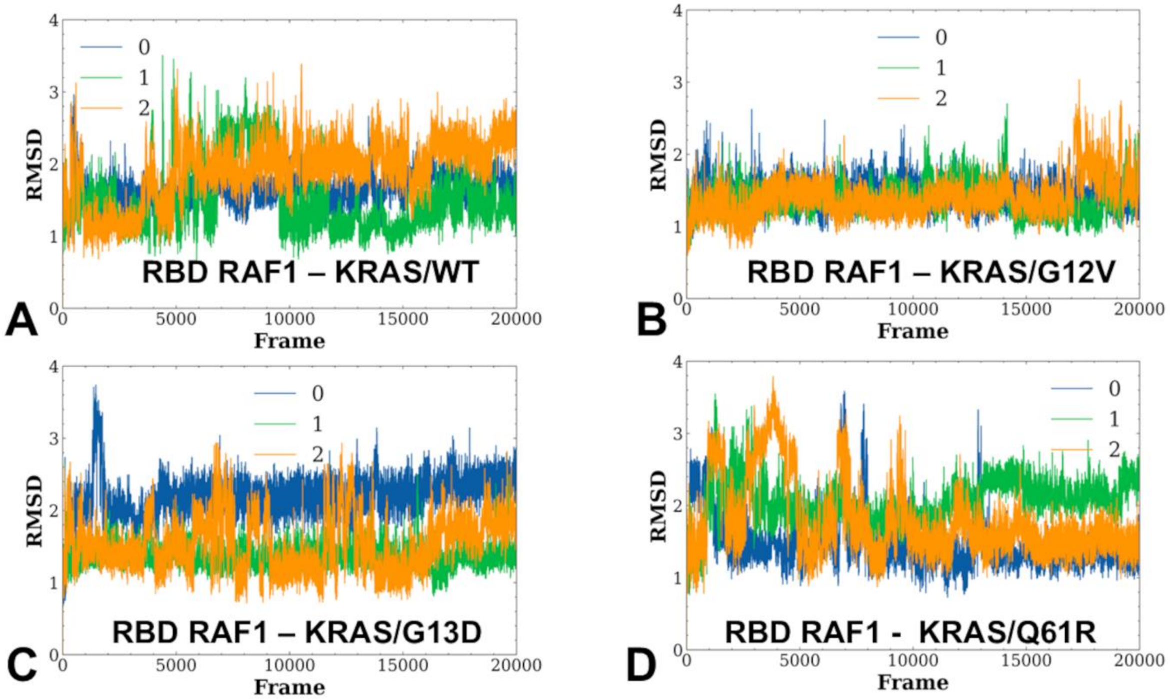


**Figure S2.** Conformational dynamics profiles of the RBD RAF1 protein residues obtained from three independent MD simulations of the KRAS complexes with RAF1. The RMSD profiles for the RBD residues obtained from three independent 2 μs MD simulations of the crystal structure of wild-type KRAS (GMPPNP-bound) in complex with the RBD of RAF1/CRAF (pdb id 6VJJ); (A), KRAS-G12V mutant in the complex with the RBD of RAF1/CRAF (pdb id 6VJJ); (B), crystal structure of KRAS-G13D (GMPPNP-bound) in complex with the RBD and CRD of RAF1/CRAF (pdb id 6XGV) (C) and the crystal structure of KRAS-Q61R (GMPPNP-bound) in complex with RBD and CRD of RAF1/CRAF (pdb id 6XGU) (D).

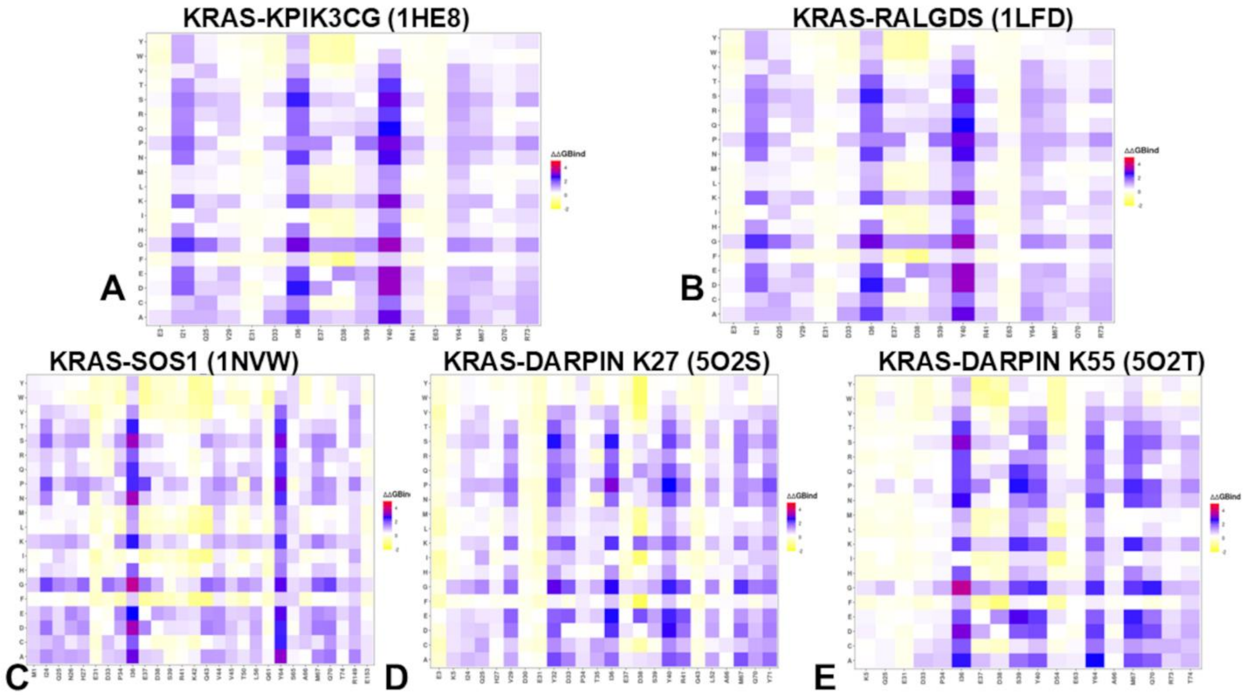


**Figure S3.** Structure-based mutational profiling of the KRAS intermolecular interfaces in the KRAS complexes with KPIK3CG (A), RALGDS (B), SOS1 (C), DARPin K27 (D), and DARPin K55 (E) . The heatmaps show the computed binding free energy changes for 20 single mutations of the interfacial positions.

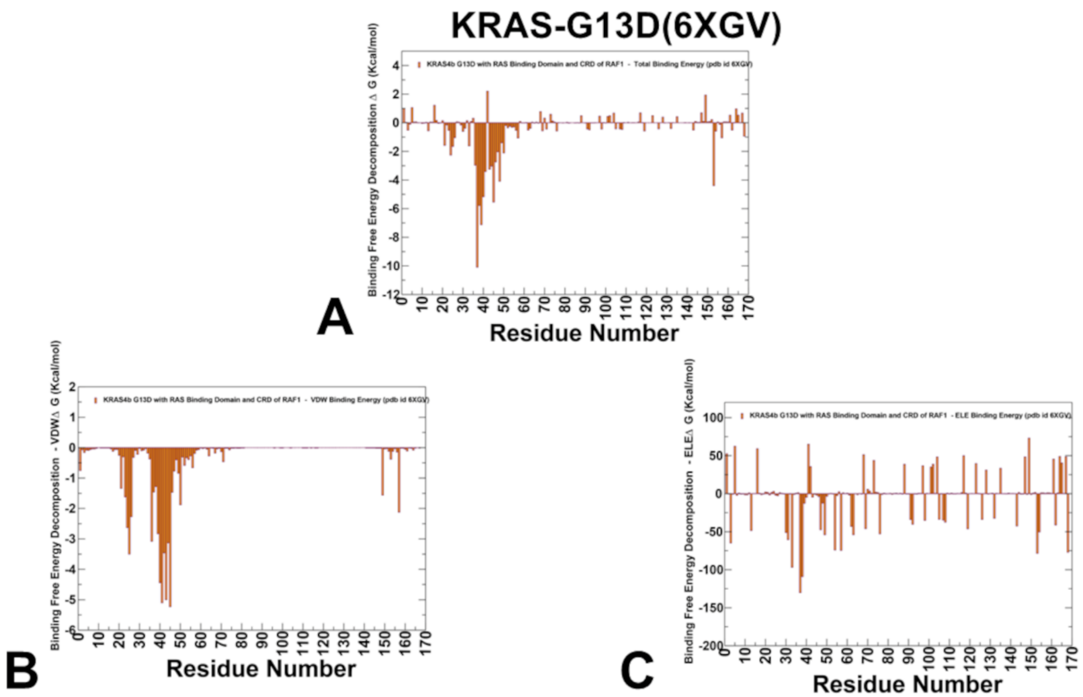


**Figure S4.** The residue-based decomposition of the total binding MM-GBSA energies for the KRAS residues in the KRAS-G13D mutant complex with RBD-CRD of RAF1 (pdb id 6XGV) (A). The residue-based decomposition of the van der Waals contribution to the total MM-GBSA binding energy for the KRAS residues in the KRAS-G13D mutant complex with RBD-CRD of RAF1 (B) and the residue-based decomposition of the electrostatic contribution to the total MM-GBSA binding energy for the KRAS-G13D mutant complex with RBD-CRD of RAF1. The MM-GBSA contributions are evaluated using 1,000 samples from the equilibrium MD simulations of respective KRAS-RAF1 complexes.

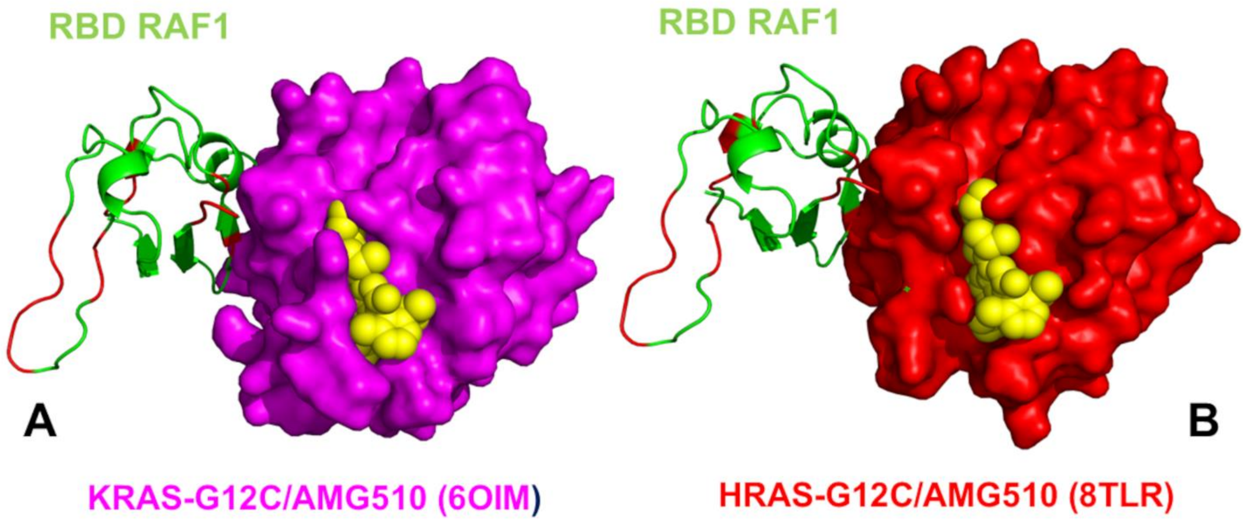


**Figure S5.** (A) The crystal structure of the human KRAS-G12C complex with RAF1 and allosteric inhibitor AMG510 KRAS-G12C is in magenta surface. RBD RAF1 is in green ribbons. AMG510 is in yellow spheres. The crystal structure of the human KRAS-G12C complex with RAF1 and allosteric inhibitor AMG510. (B) The crystal structure of the HRAS-G12C complex with RAF1 and allosteric inhibitor AMG510. HRAS-G12C complex (red surface), RBD RAF1 is in green ribbons and bound allosteric inhibitor AMG510 is yellow spheres.
